# Efficient megakaryopoiesis and platelet production require phospholipid remodeling and PUFA uptake through CD36

**DOI:** 10.1101/2023.02.12.527706

**Authors:** Maria N Barrachina, Gerard Pernes, Isabelle C Becker, Isabelle Allaeys, Thomas I. Hirsch, Dafna J Groeneveld, Abdullah O. Khan, Daniela Freire, Karen Guo, Estelle Carminita, Pooranee K Morgan, Thomas J Collins, Natalie A Mellett, Zimu Wei, Ibrahim Almazni, Joseph E. Italiano, James Luyendyk, Peter J Meikle, Mark Puder, Neil V. Morgan, Eric Boilard, Andrew J Murphy, Kellie R Machlus

**Affiliations:** Vascular Biology Program, Boston Children’s Hospital, Boston, MA, 02115 USA; Harvard Medical School, Department of Surgery, Boston Children’s Hospital, Boston, MA, 02115 USA; Haematopoiesis and Leukocyte Biology, Baker Heart and Diabetes Institute, Melbourne, VIC, Australia; Centre de Recherche du CHU de Québec-Université Laval and Centre de Recherche ARThrite, Québec, QC, G1V4G2 Canada; Department of Pathobiology and Diagnostic Investigation, Michigan State University, East Lansing, MI, USA; Institute of Cardiovascular Sciences, College of Medical and Dental Sciences, University of Birmingham, Vincent Drive, Birmingham, U.K, B15 2TT; MRC Weatherall Institute of Molecular Medicine, Radcliffe Department of Medicine and National Institute of Health Research (NIHR) Oxford Biomedical Research Centre, University of Oxford, Oxford, U.K. OX3 9DS; Metabolomics, Baker Heart and Diabetes Institute, Melbourne, VIC, Australia

**Author notes:** <u>Correspondence to:</u> Kellie R Machlus, 1 Blackfan Circle, Karp 11.214, Boston, MA 02115.

## Abstract

Lipids contribute to hematopoiesis and membrane properties and dynamics, however, little is known about the role of lipids in megakaryopoiesis. Here, a lipidomic analysis of megakaryocyte progenitors, megakaryocytes, and platelets revealed a unique lipidome progressively enriched in polyunsaturated fatty acid (PUFA)-containing phospholipids. In vitro, inhibition of both exogenous fatty acid functionalization and uptake and de novo lipogenesis impaired megakaryocyte differentiation and proplatelet production. In vivo, mice on a high saturated fatty acid diet had significantly lower platelet counts, which was prevented by eating a PUFA-enriched diet. Fatty acid uptake was largely dependent on CD36, and its deletion in mice resulted in thrombocytopenia. Moreover, patients with a CD36 loss-of-function mutation exhibited thrombocytopenia and increased bleeding. Our results suggest that fatty acid uptake and regulation is essential for megakaryocyte maturation and platelet production, and that changes in dietary fatty acids may be a novel and viable target to modulate platelet counts.

---

Lipids are key for many cell biological processes including membrane structuring, organelle compartmentalization, energy storage, and the assembly of signaling effectors.^1^ In addition, recent studies have demonstrated a role for lipids in cell fate decisions during hematopoiesis.^2–5^ Cells obtain lipids in several ways; essential fatty acids are taken up from the diet (mostly polyunsaturated fatty acids (PUFAs, containing multiple double bonds), while saturated fatty acids (SFAs, no double bonds) can be produced via *de novo* lipogenesis.^6^ As such, how lipids are produced and incorporated into cells can be regulated through cellular metabolism and dietary intervention.^5^ Megakaryocytes (MKs) are large hematopoietic cells that primarily reside in the bone marrow and produce platelets, which are essential for hemostasis.^7, 8^ Even though MKs have an extensive, lipid-rich membrane system, the role of lipids in their maturation and during platelet production has not been extensively investigated.

Megakaryopoiesis is the process by which MKs develop from hematopoietic stem cells (HSCs) along the myeloid branch of hematopoiesis under the direction of thrombopoietin (TPO) signaling through its receptor MPL. According to the classical model, each mature MK is derived from an HSC that sequentially transitions through the multipotent progenitor (MPP), common myeloid progenitor (CMP), MK-erythroid progenitor (MEP), and MK progenitor (MKP) state.^9^ After MKs are terminally differentiated, they undergo maturation^7, 8^ which includes increasing in size and developing an extensive demarcation membrane system (DMS). The DMS is a highly intertwined membrane network with numerous side branches and multiple connections with the cell surface which serves as the membrane reservoir for proplatelet formation and ultimately becomes the plasma membrane of platelets.^8, 10^ MKs then extend long proplatelet extensions through endothelial cells and into the vessel lumen and bloodstream, where they rapidly undergo repeated rounds of fission, becoming 1-3 μm circulating platelets.^7, 8^ The identification of modulators of MK maturation and platelet production is essential, as thrombocytopenia (platelet counts < 150×10^9^/L) can be life-threatening due to a heightened risk of bleeding.^11, 12^ Current standard of care is limited to therapeutics such as TPO receptor agonists which can have severe side effects including bone marrow fibrosis and leukemic transformation.^13–15^ Therefore, there is an urgent need to identify new thrombopoietic agents to increase platelet counts.

Due to the extensive DMS unique to MKs, we postulated that MKs may be more reliant on a particular membrane lipid composition than other cell types. Specifically, the processes of DMS formation and proplatelet production require a profound reorganization of both the MK cytoskeleton and the accompanying membrane system as the DMS folds and then extrudes itself outward and subsequently thins into proplatelet shafts.^7, 8^ To accomplish this, the MK membrane must acquire the lipids necessary over the course of its maturation to have sufficient flexibility for these processes. Critically, the higher numbers of double bonds in PUFAs significantly enhance membrane fluidity.^16^

While the function of lipids in platelet production remains ambiguous, recent work has begun to suggest a role for lipids in MK maturation. Valet et al.^17^ showed that MKs can take up fatty acids released by adipocytes via CD36 to facilitate their maturation *in vitro*. In addition, Kelly et al. demonstrated that the *de novo* lipogenesis pathway can regulate late-stage MK maturation and platelet formation.^18^ These studies support a role for lipids in MK maturation and raise further questions about the relationship between megakaryopoiesis and lipid biosynthesis. Here, we expand the previously suggested role of lipids in MKs by performing lipidomics to uncover the lipid fingerprint of MEPs, immature and mature MKs, and platelets. We demonstrate that altering both *de novo* lipogenesis and fatty acid functionalization and uptake abrogate megakaryopoiesis and proplatelet formation. Further, we reveal that platelet counts can be modulated *in vivo* by altering dietary fatty acid content. Finally, we identify CD36 as a key fatty acid uptake receptor that affects platelet counts in both mice and humans. These data support a key role for fatty acids in MK maturation and platelet production and suggest that dietary interventions can influence thrombopoiesis.

## RESULTS

### Megakaryocytes and platelets display a unique phospholipid profile enriched in polyunsaturated fatty acids

To provide insight into how the lipid profile changes throughout MK differentiation and maturation, we performed an extensive lipidomic study of MK maturation using liquid chromatography tandem-mass spectrometry (LC-MS/MS) starting with MEPs. The indicated cell populations were sorted from adult murine bone marrow and platelets were isolated from autologous blood (Fig. 1A). Dimensionality reduction of all populations revealed that MKs and platelets have a lipidome distinct from their precursor, MEPs (Fig. 1B), which was confirmed when analyzing different lipid class compositions (Supplementary Fig. S1A). These data suggest that extensive remodeling of the lipidome occurs during megakaryopoiesis. Further, when comparing the lipid composition of the microenvironment to the cells that reside in it, such as comparing bone marrow extracellular fluid (BMEF) to bone marrow cells (Supplementary Fig. S1B-D) or plasma to platelets (Fig. S1E), we found that the lipid composition of these cells (Supplementary Fig. SF-I) was unique to their environments. We used a lipid ontology analysis to identify the main lipid species that varied between the different cell populations and found that membrane lipids, glycerophospholipids, and fatty acids were upregulated over the course of megakaryo- and thrombopoiesis (Fig. 1C). This was further supported by analysis of an mRNA sequencing dataset previously published by our group^19^ which revealed that MKs are actively regulating key lipid-related mRNAs as they undergo proplatelet formation (Fig. 1D-E), specifically highlighting pathways involved in fatty acid metabolism and uptake. A pathway analysis further emphasized the synthesis of the long chain fatty acyl-CoA and fatty acid metabolism as key pathways involved in proplatelet production (Fig. 1E). Therefore, as lipidomic and mRNA pathway analyses suggested that differences in phospholipids and fatty acids were unique and important to MK maturation and platelet production, we analyzed these lipid classes in our cell populations. Our data revealed that MKs and platelets were enriched in phosphatidylcholine (PC) compared to their progenitors. Conversely, MKs and platelets were reduced in phosphatidylethanolamine (PE), phosphatidylglycerol (PG), and phosphatidylinositol (PI) (Fig. 1F). When examining the differences in the overall fatty acid saturation level, we identified a significant reduction in saturated fatty acids (no double bonds) along the maturation pathway (Fig 1G-H). Notably, we also found that cells acquired increasing levels of more complex PUFA-containing phospholipids (6+ double bonds, Fig 1G, I) as they matured, with platelets exhibiting the highest levels. This overall pattern of fatty acid remodeling was also seen in other phospholipid classes such as PC, PE, PI and phosphatidylserine (PS) (Supplementary Fig. S1 D-G). Taken together, these data reveal significant phospholipid remodeling, and specifically an increase in PUFAs during megakaryopoiesis.

**Figure 1.**
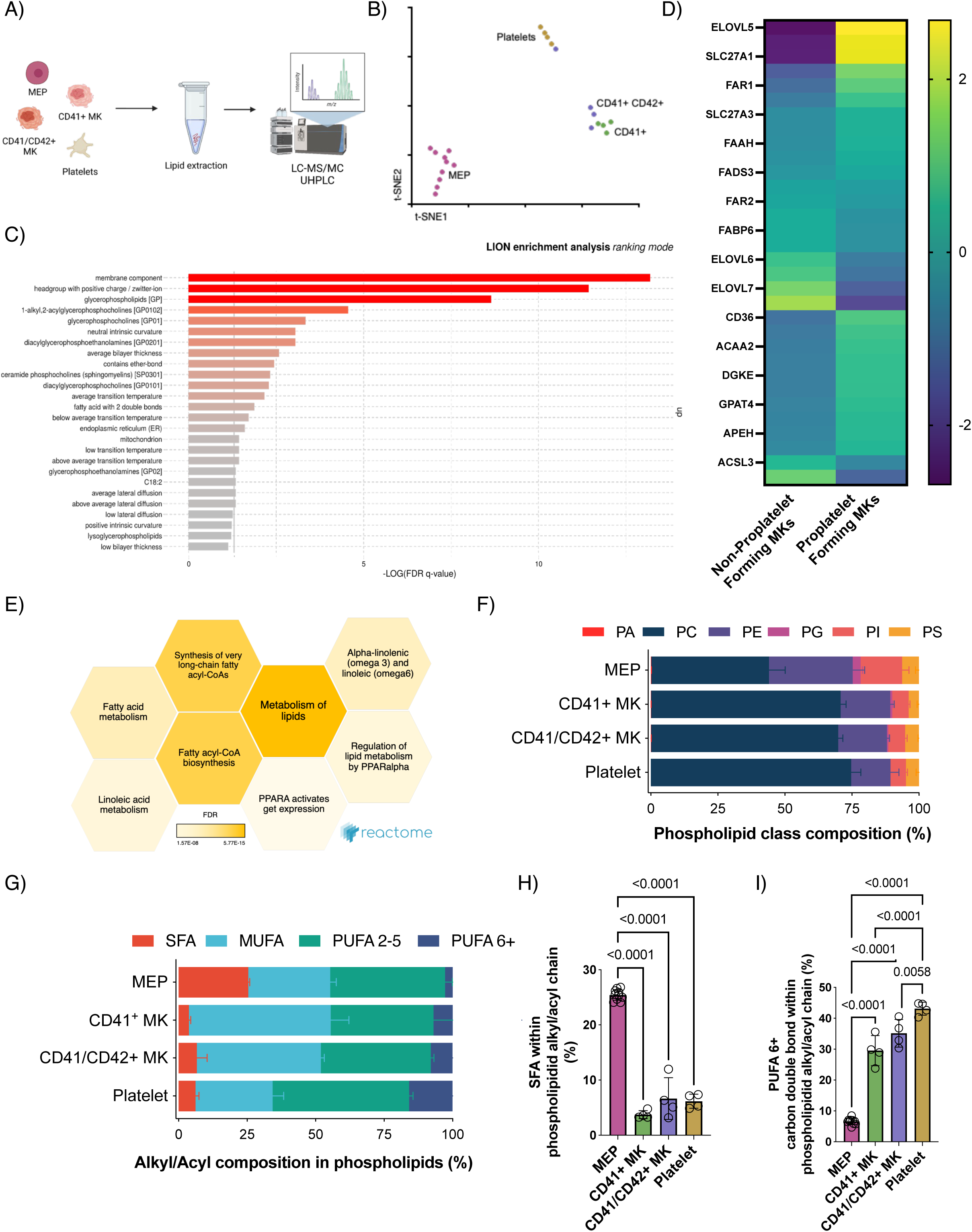
Megakaryocytes and platelets have a unique lipid profile that is enriched in polyunsaturated fatty acids. (A) Murine bone marrow cell populations were isolated by fluorescence-activated cell sorting and platelets by sequential centrifugation. Lipids were extracted and analyzed using 20-min gradient HPLC and mass spectrometry (see methods for details), n=8 for MEP and n=4 for all other cell populations. (B) T-distributed stochastic neighbor embedding analysis (tSNE) highlights lipidomic differences between MEPs, immature (CD41+) MKs, mature (CD41/42+) MKs, and platelets. (C) LION enrichment analysis showing the top 10 upregulated LION terms. (D) Bulk RNA sequencing was performed on MKs immediately preceding and during proplatelet formation^19^ and heatmap shows the log2FoldChange of selected genes. (E) Reactome enrichment analysis from bulk RNA sequencing on MKs reveals that canonical pathways altered include metabolism of lipids, synthesis of very long-chain fatty acyl-CoAs, and fatty acyl-CoA biosynthesis. Color intensity is directly correlated to the false discovery rate. Total percentage of (F) phospholipid classes and (G) lipid saturation levels of indicated murine bone marrow cell populations in lipidomics analyses. Percentage of SFAs (H) and PUFAs with 6+ double bonds (I) in indicated cell populations. Illustrations were done with Biorender® *MK: megakaryocyte; MEP: MK-erythroid progenitor; PA: phosphatidic acid; PC: phosphatidylcholine; PE: phosphatidylethanolamine; PI: phosphatidylinositol; PS: phosphatidylserine; PG: phosphatidylglycerol; SFA: saturated fatty acid; MUFA: monounsaturated fatty acid; PUFA: polyunsaturated fatty acid*

### Manipulation of fatty acid synthesis impairs megakaryocyte maturation and proplatelet formation

Since we identified that PUFA content increases throughout megakaryopoiesis, we hypothesized that PUFA uptake is important for MK maturation and platelet production. PUFAs are either considered essential fatty acids and are therefore obtained from the diet, or are synthesized from essential fatty acid precursors. To test the importance of fatty acid uptake and functionalization in MK differentiation and proplatelet formation, we inhibited ACSL, the enzyme that catalyzes the formation of acyl-CoA from fatty acids, a necessary step for the incorporation of fatty acids into phospholipids (Fig. 2A). We isolated murine bone marrow- (Fig. 2) and fetal liver-derived (Supplementary Fig. S2) hematopoietic stem and progenitor cells (HSPCs) and cultured them with TPO for 4 days with the indicated concentrations of Triacsin C. Inhibition of ACSL led to a significant, dose dependent reduction in the number of immature (CD41^+^) and mature (CD41/42^+^) MKs in both bone marrow- (Fig. 2B-C) and fetal liver(Supplementary Fig. S2 A)-derived MKs.

**Figure 2.**
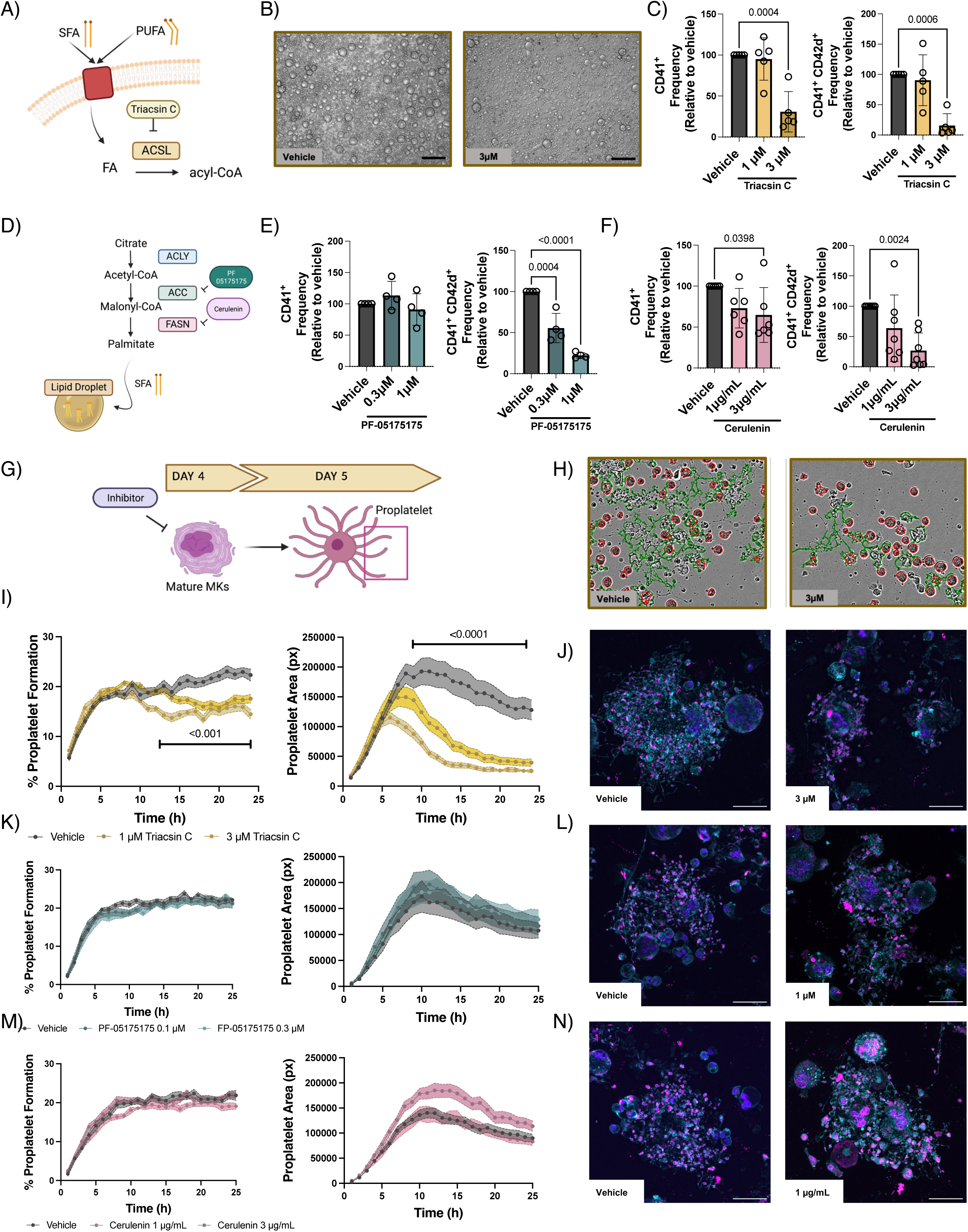
Fatty acid incorporation and de novo lipogenesis are necessary for MK differentiation and efficient proplatelet formation. (A) Schematic of fatty acid functionalization through long chain acyl-coA synthetase (ACSL). Murine bone marrow HSPCs were cultured with TPO and treated with the ACSL inhibitor Triacsin C (1 M and 3 M). (B) Representative images after 4 days of treatment, showing MKs (large cells) and surrounding HSPCs, which failed to differentiate. n=5, scale bar=150 m (C) CD41+ and CD41/CD42d+ cells were quantified using flow cytometry, n=5, one-way ANOVA – Dunnett’s test (D) Schematic of *de novo* lipogenesis. Murine bone marrow HSPCs were cultured with TPO and treated on day 0 with the acetyl-CoA carboxylase (ACC) and fatty acid synthetase (FASN) inhibitors, (E) PF-05175157 (0.3 and 1 M) n=4 and (F) Cerulenin (1 and 3 g/mL) n=6, respectively. CD41+ and CD41/42d+ positive cells were quantified by flow cytometry, one-way ANOVA – Dunnett’s test. (G) To quantify effects on proplatelet formation, mature fetal liver MKs (day 4) were treated with inhibitors at indicated dosages and proplatelet percentage and area were quantified using the Incucyte high content imaging system. (H) Representative phase contrast image after treating mature FLMKs with Triacsin C showing example quantification of round (red outline) versus proplatelet-making (green outline) MKs (vehicle, left, and Triacsin C, 3 M, right). (I) Representative graph of proplatelet formation and proplatelet area after treating FLMKs with vehicle (grey) and Triacsin C (1 and 3 M, dark and light yellow, respectively), n=3, Two-way Anova. (J) Representative images of bone marrow MKs using confocal microscopy after treating mature MKs for 24 hours (vehicle left, Triacsin 3 M, right). b-tubulin (cyan), phalloidin (magenta), DAPI (blue). Scale bar=50 m (K) Representative graph of proplatelet formation and proplatelet area after treating FLMKs with vehicle (grey) and ACC inhibitor, PF-05175175 (0.1 and 0.3 M, dark and light green, respectively), n=3, Two-way Anova. L) Representative images of bone marrow MKs using confocal microscopy after treating mature MKs for 24 hours (vehicle left, PF-05175175 1 M, right). b-tubulin (cyan), phalloidin (magenta), DAPI (blue). Scale bar=50 m. (M) Representative graph of proplatelet formation and proplatelet area after treating FLMKs with vehicle (grey) and FASN inhibitor, cerulenin (1 and 3 g/mL, dark and light pink, respectively), n=3, Two-way Anova. N) Representative images of bone marrow MKs using confocal microscopy after treating mature MKs for 24 hours (vehicle left, cerulenin 1 g/mL, right). b-tubulin (cyan), phalloidin (magenta), DAPI (blue). Scale bar=50 m. Illustrations were done with Biorender®.

Next, we inhibited *de novo* lipogenesis using inhibitors of acyl-coA carboxylase (ACC) (PF-05175175) and fatty acid synthetase (FASN) (Cerulenin) (Fig. 2D). Both inhibitors significantly and dose dependently reduced MK maturation in bone marrow- (Fig. 2E-F) and fetal liver-derived (Supplementary Fig. S2 C, E) derived MKs. Notably, the frequency of mature MKs (CD41/42^+^) was decreased more than immature MKs (CD41^+^). None of the inhibitors were cytotoxic (Supplementary Fig. S2 B, D, F), suggesting that the effects on HSPCs were due to a failure in MK differentiation.

Once mature, MKs remodel the DMS into proplatelets. We postulated that this process is dependent on a highly specific membrane lipid content to allow for proplatelet elaboration. To explore the role of fatty acid uptake and synthesis on proplatelet formation, we treated mature MKs (day 4 of culture, i.e. 24h preceding proplatelet formation) with the indicated inhibitors and monitored proplatelet formation over 24 hours (Fig. 2G). Triacsin C treatment resulted in a significant reduction in both the number of MKs making proplatelets and the area of formed proplatelets (Fig. 2H-J), suggesting a severe impairment in proplatelet elaboration. However, neither of the *de novo* lipogenesis inhibitors significantly impacted proplatelet formation (Fig. 2K-N), indicating that *de novo* lipogenesis may not be essential for proplatelet generation. To exclude the possibility that effects on proplatelet formation were due to impaired mitochondrial activity, we measured oxygen consumption rate using a Seahorse mitostress assay of murine bone marrow-derived MKs treated with the indicated inhibitors. The MK mitochondrial profile was not altered upon inhibition of fatty acid synthesis with Triacsin C, Cerulenin, or PF-051751 (Supplementary Fig. S2 G-H), supporting our conclusion that the observed changes in MK maturation and/or proplatelet formation were due to a role of fatty acid incorporation or synthesis and not mitochondrial metabolism. Together, these data suggest that MK differentiation and maturation are reliant on both fatty acid uptake and *de novo* lipogenesis. However, proplatelet formation appears uniquely reliant on fatty acid uptake and functionalization.

### Administration of a high saturated fatty acid diet increases megakaryocyte size and reduces platelet count

Our data demonstrate that megakaryopoiesis and platelet production are dependent on fatty acid uptake, functionalization, and metabolism. We therefore postulated that altering the exogenous supply of fatty acids will alter MK phenotype and subsequent platelet production. We first wanted to examine the effects of direct supplementation of SFAs on MK development and platelet production. To confirm and visualize SFA incorporation into MKs in vitro, we performed click-chemistry using palmitic acid modified with a terminal alkyne group. First, murine bone marrow HSPCs were supplemented with modified SFA in culture. After 4 days, mature MKs were functionalized with an azide-linked fluorescent reporter, which bound modified palmitic acid in the MK membrane (Fig. 3A), allowing visualization (Fig. 3B). We observed a robust, dose-dependent incorporation of palmitic acid throughout the plasma and demarcation membrane of MKs (Fig. 3B-C). Additionally, MKs supplemented with palmitic acid were significantly larger and displayed a reduced capacity to form proplatelets in vitro (Fig. 3D-E). These data confirmed that MKs incorporated SFAs (palmitic acid) into their membrane as they differentiate and mature, resulting in increased size and decreased proplatelet generation.

**Figure 3.**
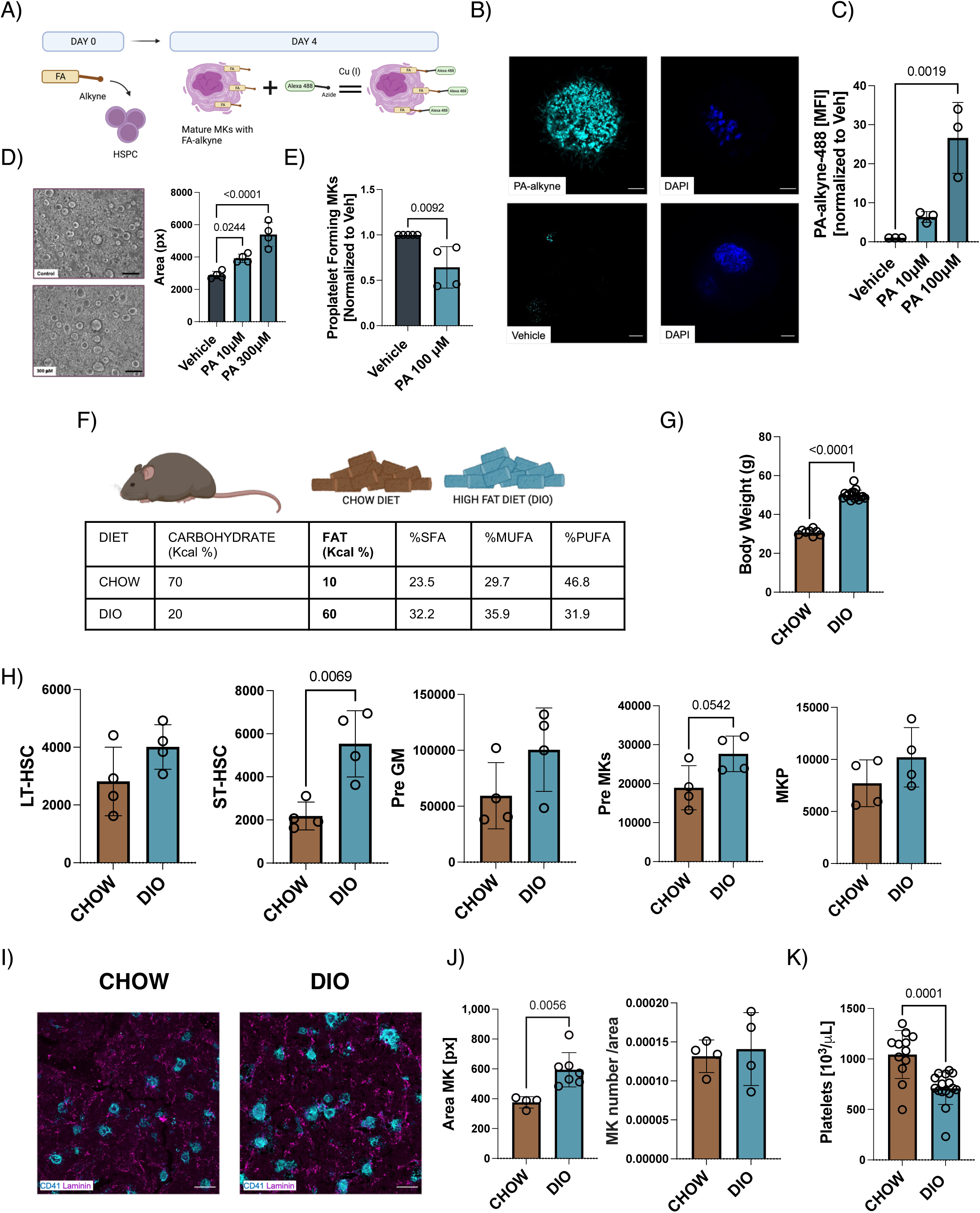
A high saturated fat diet significantly alters MK phenotype and reduces platelet counts. (A) Schematic of click-chemistry technique where fatty acids modified with an alkyne group are supplemented into cultured murine bone marrow HSPCs. After 4 days of culture, the incorporated alkyne in mature MKs was functionalized with an azide-linked fluorescent reporter to visualize fatty acid incorporation into cells. (B) Murine bone marrow HSPCs from wildtype mice were incubated with 100 M of modified palmitic acid. After 4 days of maturation, MKs were isolated, and the azide-linked fluorescent reported was added. The incorporation of palmitic acid (PA) was visualized using confocal microscopy. PA-alkyne (cyan); DAPI (blue). Scale bar=5 m (C) MFI of MKs incubated with 10 and 100 M PA was calculated using ImageJ. n=3, one-way ANOVA – Dunnett’s test (D) Cultured murine bone marrow HSPCs from wildtype mice were cultured with TPO and supplemented with 10 and 300 M palmitic acid. Representative images show MK size. MK area was quantified using ImageJ. n=4, one-way ANOVA – Dunnett’s test, scale bar=150 m. (E) To examine proplatelet formation, mature MKs (day 4) were supplemented with palmitic acid (100 M) or DMEM with 0.1% BSA (vehicle), the percentage of MKs making proplatelets at 24h was quantified using the Incucyte high content imaging system. n=4, unpaired t-test (F) Male mice were fed a 60% high fat (D12492, Research Diets Inc) or chow diet (D12450B, Research Diets Inc) for 14 weeks. n=16, unpaired t-test (G) Mice were weighed at week 14, n=16 mice per group (H) LT-HSC, ST-HSC, Pre MKs, and MKP cell populations were quantified at week 14 using flow cytometry, n=4, unpaired t-test (I) Representative images of bone marrow showing MKs (CD41, blue) and vasculature (laminin, pink) in femur cryosections. Scale bar=50 m (J) MK area and number were quantified manually from femur cryosections using ImageJ. n=4-7, unpaired t-test (K) Platelet counts were measured using a Sysmex hematology analyzer. n=12-16, unpaired t-test. Illustrations were done with Biorender®.

To test the impact of an SFA-enriched diet in vivo, male mice were fed a 60% high fat diet (“diet-induced obesity (DIO) model”) for 14 weeks (Fig. 3F), which led to increased body weight in the DIO mice (Fig. 3G). To determine whether the high fat diet affected HSPC differentiation, we quantified the number of bone marrow HSPCs. While we did not detect differences between long term (LT)-HSC, Pre-GM, or Pre-MK populations after 14 weeks, we found a significant increase in short-term (ST)-HSCs and Pre-MKs (Fig. 3H). Further, bone marrow MKs were significantly larger in DIO mice (Fig. 3I-J), consistent with our in vitro data (Fig. 3D), while their numbers were unchanged. Finally, platelet counts were significantly reduced after administration of the SFA-enriched diet (Fig. 3K). No significant differences were found in other blood parameters (Supplementary Fig. S3 A-E). These data revealed that enriched dietary SFAs increased MK size and decreased platelet production both in vitro and in vivo. Further, these data support our hypothesis that enhanced membrane PUFA content is necessary for maximal proplatelet production, and that interruption of this process disrupts platelet production.

### Administering a high polyunsaturated fatty acid ratio in a high fat diet prevents a reduction in platelet counts

Feeding mice a high fat diet with an enrichment in SFA resulted in decreased platelet counts. This is consistent with our lipidomic data, which revealed a clear bias towards PUFAs, and not SFAs, during thrombopoiesis. Thus, we hypothesized that maintaining a higher proportion of PUFAs in the high fat diet may prevent obese mice from having reduced platelet counts. To explore this, we first establish whether MKs could take up PUFAs, and examined their effect *in vitro* by performing click chemistry as described above. Indeed, MKs dose-dependently incorporated arachidonic acid (Fig. 4A), however it did not significantly alter their area (Fig. 4B) or capacity to form proplatelets *in vitro* (Fig. 4C).

**Figure 4.**
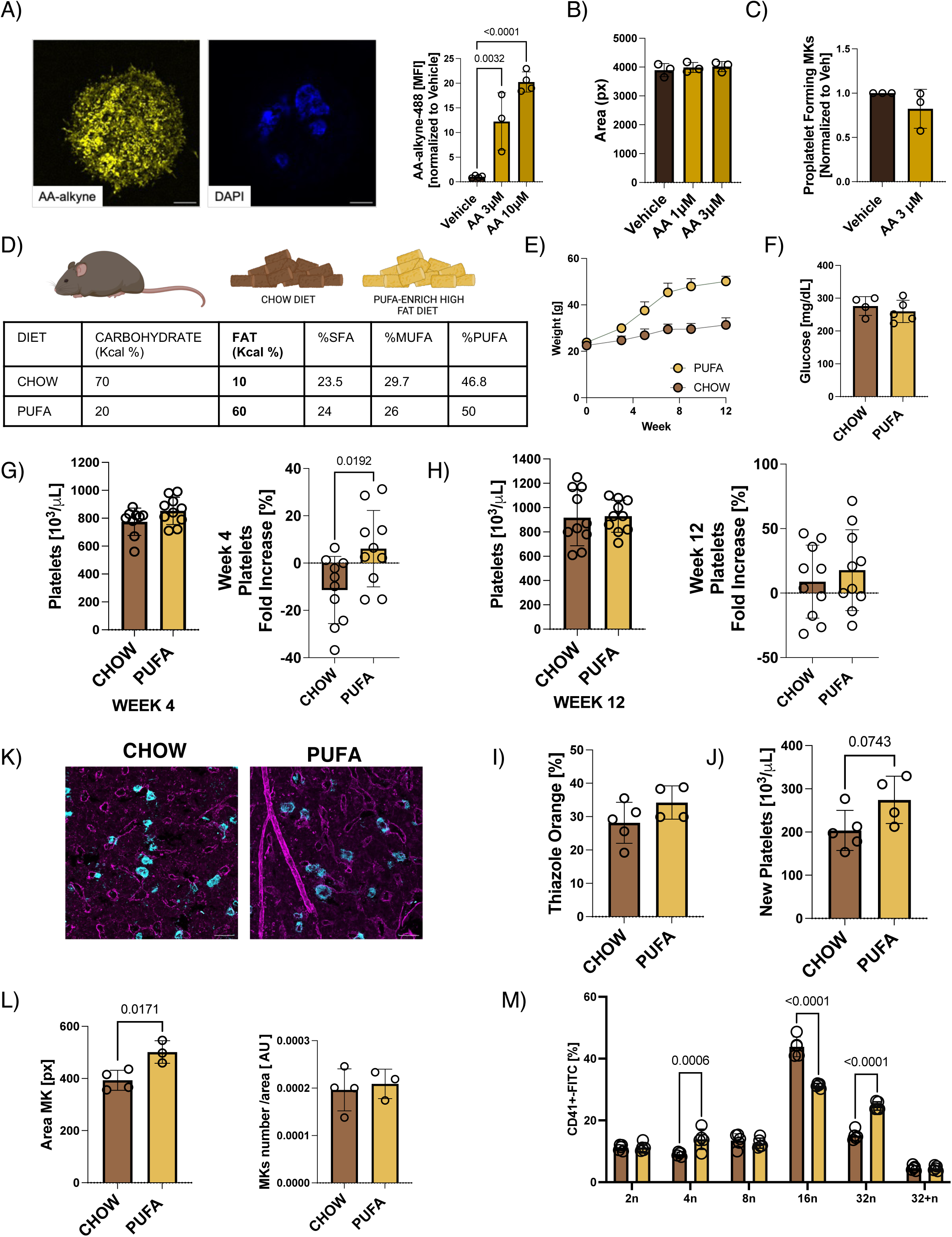
Platelet counts can be modified in vivo by altering dietary polyunsaturated fatty acid composition. (A) Murine bone marrow HSPCs were incubated with arachidonic acid modified with an alkyne group (3 and 10 M). After 4 days of maturation, MKs were isolated and a fluorescently-conjugated azide was added. Incorporation of arachidonic acid was visualized using confocal microscopy and MFI was calculated using ImageJ. n=3, one-way ANOVA – Dunnett’s test. Yellow: AA-alkyne (yellow), DAPI (blue); Scale bar=5 m. (B) Murine bone marrow HSPCs from wildtype mice were cultured with TPO and media supplemented with arachidonic acid (3 and 10 M). MK area was quantified using ImageJ. n=6 and n=3, respectively, n=3, one- way ANOVA – Dunnett’s test (C) Mature MKs (day 4) were cultured arachidonic acid at indicated dosages and the percentage of MKs making proplatelets at 24h was imaged using the Incucyte high content imaging system. n=3, unpaired t-test (D) Male mice were fed a 60% high fat diet enriched in polyunsaturated fatty acids (PUFAs) (D22050406i, Research Diets Inc) or chow diet (D12450B, Research Diets Inc) for 12 weeks. n=10 mice per group. Body weight (E) and glucose levels (F) were measured, n=10, unpaired t-test. Platelet counts were measured at week 4 (G) and 12 (H) using a Sysmex hematology analyzer. n=10, unpaired t-test. Newly made platelets were analyzed by quantifying (I) percentage and (J) absolute platelet numbers positive for thiazole orange by flow cytometry, n=4-5, unpaired t-test. (K) Representative images showing MKs (CD41, blue) and vasculature (laminin, pink) in femur cryosections at week 12. (L) MK area and number were quantified from femur cryosections using ImageJ, n=3-4, unpaired t-test. (M) Ploidy analysis of native bone marrow MKs assessed by propidium iodide staining and quantified by flow cytometry. n=5, two-way ANOVA. Illustrations were done with Biorender®.

To determine whether the low platelet counts observed in the DIO model were a consequence of the high fat diet or the fatty acid saturation status (high SFA), we fed male mice a matched 60% high fat diet enriched in PUFAs (Fig. 4D) instead of SFAs. Mice fed the PUFA diet weighed significantly more than controls (Fig. 4E), but glucose levels were indistinguishable (Fig. 4F), suggesting the mice were not diabetic. Mice on the high PUFA diet had significantly increased platelet counts after 4 weeks, and platelet counts remained elevated at the study endpoint (Fig. 4G-H). No differences were found in other blood parameters (Supplementary Fig. S4 A-C).

In addition, when assessing the amount of circulating, newly generated reticulated platelets using thiazole orange, we found that mice fed the enriched PUFA diet had a tendency toward more immature platelets in circulation (Fig. 4I-J). Comparable to the DIO mice, MKs in the PUFA-fed mice were significantly larger than the chow group while their number remained unaltered (Fig. 4K-L). In addition, their maturation, as measured by ploidy, was substantially enhanced, with a significant decrease in the number of 16n MKs and an increase in 32n MKs (Fig. 4M), suggesting an overall shift to higher ploidy. Together, these data reveal that enhancing the amount of PUFAs in a high fat diet can rescue the reduced platelet counts seen in the DIO model. These results underscore the role of dietary PUFAs in contributing to MK maturation and reveal that supplementation with PUFAs both in vitro and in vivo can enhance MK maturation and platelet production.

### CD36 KO mice have decreased platelet counts and do not respond to high fat diets

Our data revealed that MKs and their progenitors readily take up exogenous fatty acids, and this is an essential process during their maturation. Further, modifying dietary fatty acids can directly impact platelet counts in vivo. As Valet et al. recently demonstrated that MKs can take up fatty acids via the scavenger receptor CD36 to help facilitate membrane maturation^17^, and CD36 was upregulated in proplatelet-forming MKs (Fig. 1E), we decided to explore whether CD36 is the mechanism by which MKs and their progenitors incorporated exogenous fatty acids. We utilized a mouse model constitutively lacking CD36 (*Cd36^−/−^*).^20^ *Cd36^−/−^* mice exhibited significantly reduced platelet counts (Fig. 5A).^17^ While MPV and IPF remained unchanged, red blood cell counts were also significantly reduced (Fig. 5B). To directly test if CD36-deficient MKs had a defect in taking up fatty acids in vitro, we used click-chemistry on MKs from *Cd36^−/−^* and wildtype mice cultured with either the SFA palmitic acid (Fig. 5C-D) or the PUFA arachidonic acid (Fig. 5E). In line with our hypothesis, *Cd36^−/−^* MKs took up significantly less fatty acids (Fig. 5C), suggesting that the CD36 receptor plays a substantial role in fatty acid uptake of both SFAs (Fig. 5D) and PUFAs (Fig. 5E) in MKs.

**Figure 5.**
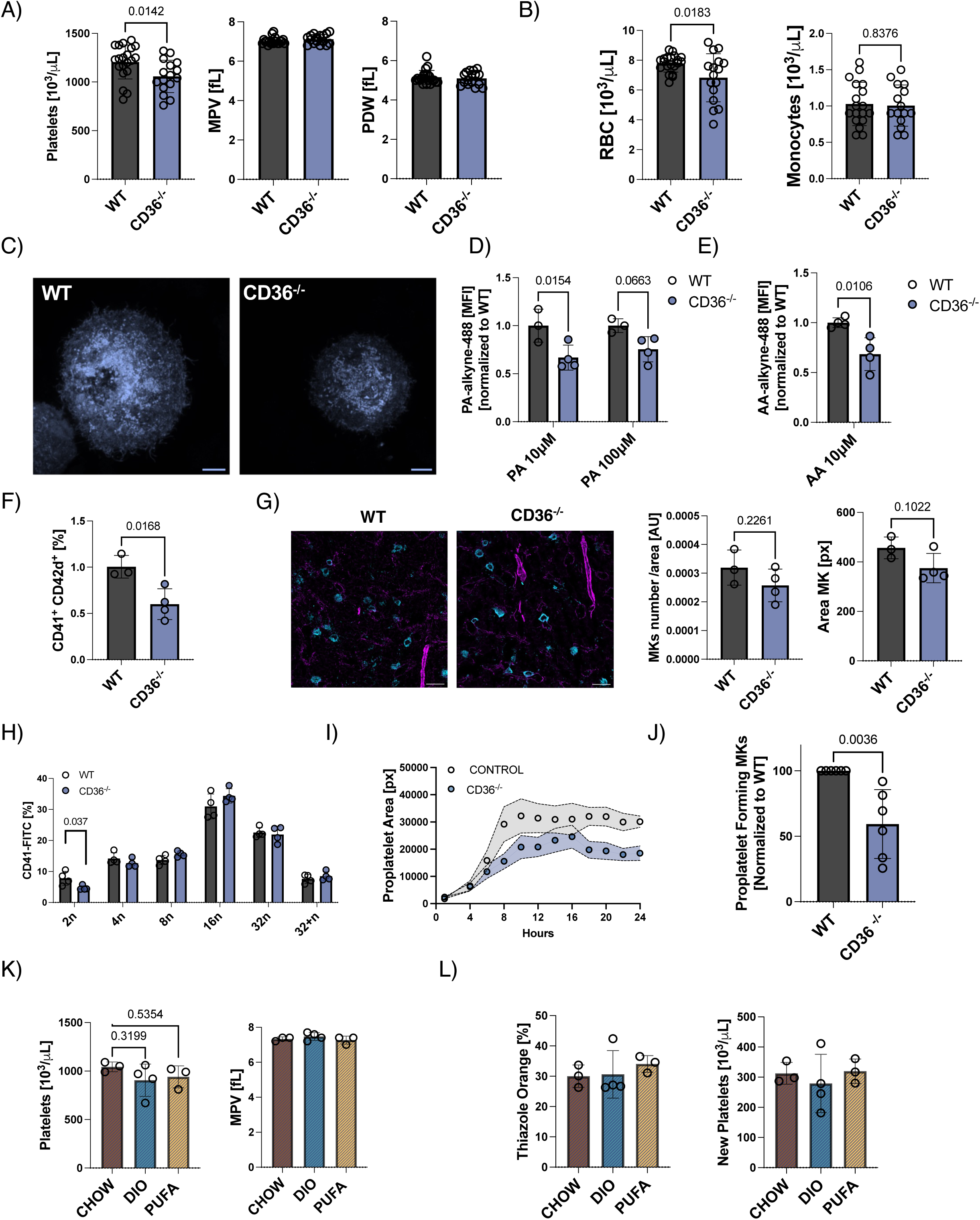
Lack of CD36 in mice reduces cellular fatty acid incorporation and impairs proplatelet formation. Platelets and megakaryocytes were characterized in adult male wildtype (WT) and *Cd36* deficient (*Cd36^−/−^*) mice. (A) Platelet counts, mean platelet volume (MPV), and platelet distribution (PDW) were measured using the Sysmex. n=16-20, unpaired t-test. (B) Red blood cell and monocyte counts were measured using Sysmex, unpaired t-test, n=16-20 (C-E) HSCs from WT and *Cd36^−/−^* mice were cultured with palmitic acid and arachidonic acid modified with an alkyne group at indicated dosages. After 4 days of maturation, MKs were isolated, and click-chemistry performed as previously described. (C) Representative images of MKs from WT and *Cd36^−/−^* mice. MFI was calculated using ImageJ. Blue: palmitic acid (100 M). scale bar=5 m. Incorporation of palmitic acid (D) and (E) arachidonic acid were visualized using confocal microscopy and MFI was calculated using ImageJ. n= 4, unpaired t-test (F) Murine bone marrow HSPCs from WT and *Cd36^−/−^* mice were cultured with TPO. After 4 days of maturation, the number of mature MKs (CD41/CD42d+ cells) was measured using flow cytometry, n=3-4, unpaired t-test. (G) MK number and area was quantified from femur cryosections using ImageJ. n=3-4, unpaired t-test. Representative images show MKs (CD41, blue) and vasculature (laminin, pink). (H) Ploidy analysis of cultured marrow MKs assessed by propidium iodide staining and flow cytometry. Percentage of CD41+ cells with different levels of ploidy is shown. n=4, two-way ANOVA. (I-J) Proplatelet formation was quantified from mature MKs (day 4) from WT and *Cd36^−/−^* mice in the presence of hirudin. (I) Representative graph of proplatelet area from n=4. (J) Representative proplatelet area, n= 4, unpaired t-test (J) and proplatelet percentage at 24h, n= 6, unpaired t-test (K) were quantified using the Incucyte high content imaging system. (K-L) Adult male WT and CD36 KO mice were fed chow, saturated- and PUFA-enriched diets as in Figures 3 and 4 for 8 weeks and K) platelet counts were measured by Sysmex. L) Newly made platelets were analyzed by quantifying percentage and absolute platelet numbers positive for thiazole orange by flow cytometry, n=3-4 mice per group, unpaired t-test.

We next cultured HSPCs derived from *Cd36^−/−^* and wildtype mice and found a significant decrease in the number of mature MKs that differentiated from *Cd36^−/−^* mice, suggesting that CD36 is important not only for platelet production but also MK differentiation (Fig. 5F). We examined the number and area of CD41+ cells in *Cd36^−/−^* bone marrow and found no significant reduction (Fig. 5G). Moreover, the ploidy of in vitro differentiated MKs was largely unchanged, with only a decrease in the 2n population (Fig. 5H). As *in vitro* MK differentiation was affected in *Cd36^−/−^* mice, we next characterized proplatelet formation. Critically, CD36-deficient MKs displayed a notable defect in proplatelet formation with both the number of MKs forming proplatelets and proplatelet area being significantly reduced (Fig. 5I-K). This finding strongly suggests that proplatelet formation is dependent on the uptake of essential fatty acids through CD36. Together, these data indicate that loss of MK CD36 decreases the uptake of both SFAs and PUFAs into MKs. Further, lack of CD36 on HSPCs decreases their ability to differentiate into MKs and the capacity of MKs to make proplatelets.

To test whether the loss of CD36 *in vivo* could disrupt the ability of both SFA- and PUFA-enriched high fat diets to modulate platelet counts, we fed mice chow or 60% high fat diets enriched in either SFAs (DIO model) or PUFAs. There were no significant differences in platelet counts (Fig. 5K), or platelet production (Fig. 5L), across the three groups after 8 weeks. This suggests that loss of CD36 abrogated the ability of dietary fatty acids to alter platelet counts *in vivo*. Together, these data further support the role of CD36 as a key receptor that takes up essential fatty acids that drive MK maturation and platelet production.

### Identification and characterization of patients with a CD36 loss of function mutation with reduced platelet counts and increased platelet volume

To investigate the biological relevance of CD36 in humans, we identified a family with an idiopathic thrombocytopenia; patients II.1 and II.2 were recruited to the UK-GAPP study (Fig. 6A), and clinical histories evaluated. Whole blood cells counts were taken, revealing low platelet counts and high MPV and IPF values (Fig. 6B, Table 1). In addition, the mother (I:2) had bleeding episodes and low platelet counts. Whole Exome Sequencing (WES) analysis was performed using a bioinformatic pipeline workflow which identified an average total of 43,884 variants. Genetic variants were filtered against a panel of 358 genes known or predicted to be associated with platelet count, function, or lifespan. These variants where then filtered out by excluding all synonymous and intronic variants followed by excluding all variants with a MAF > 0.01 (Fig. 6C). Pathogenicity prediction of the variants were determined by utilizing the prediction tools (Mutation Taster, PolyPhen-2, SIFT, Provean) and the variants were classified based on the ACMG guidelines. Plausible candidate variants in each patient were then selected based on the pathogenicity prediction (Fig. 6D). This identified a pathogenic heterozygous stop gain variant in exon 10 of the CD36 gene (c.975T>G; p. Tyr325Ter) in both patients II.1 and II.2 (Fig. 6C).

**Figure 6.**
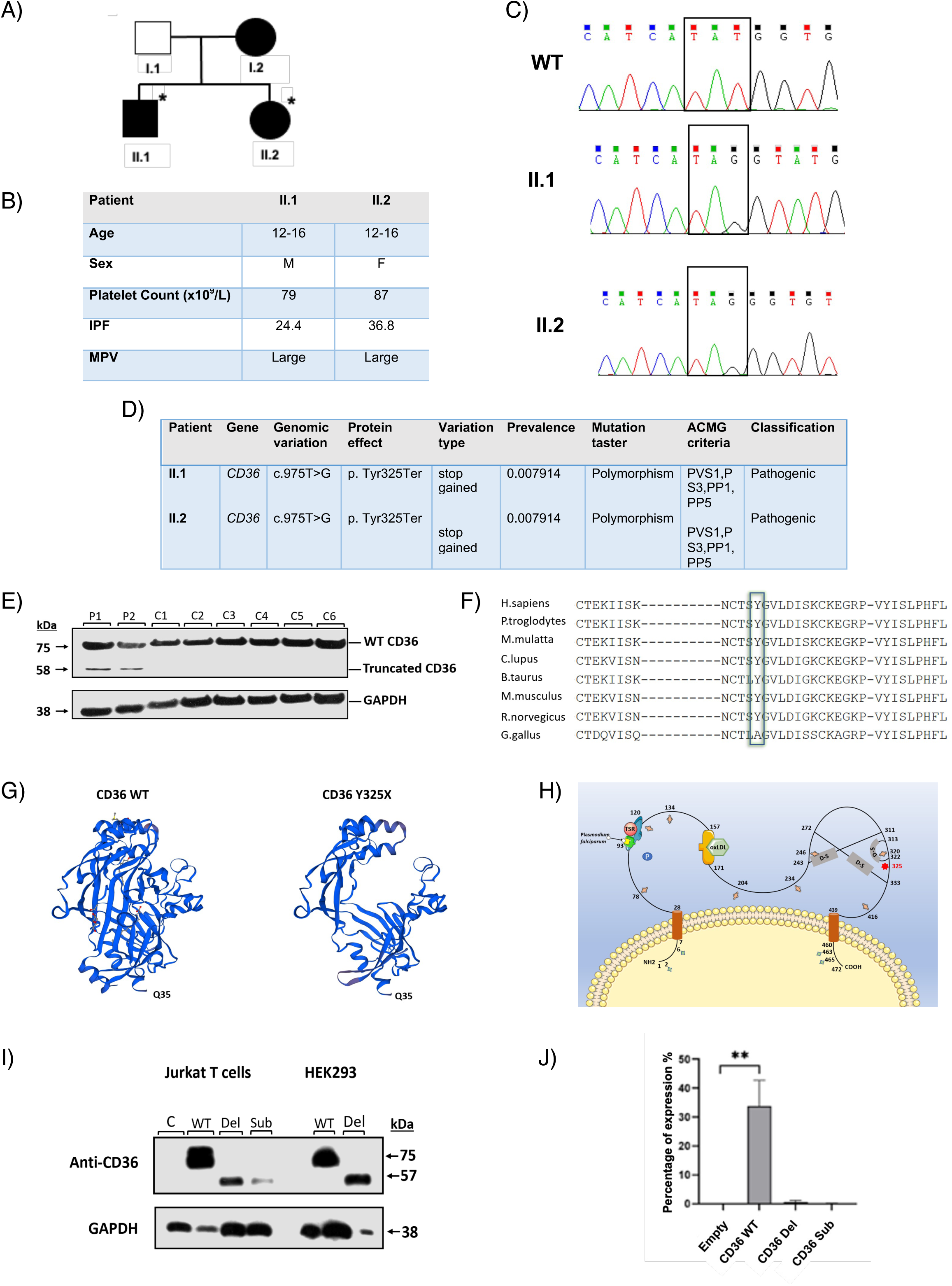
Identification of a CD36 loss-of-function variant (p.Tyr325Ter) in patients with thrombocytopenia. (A) Family pedigree including affected mother (I:2) and affected patients II.1 and II.2. Affected individuals are highlighted in solid black. The mother was reported by personal communication to have bleeding episodes and low platelet counts. Asterisks (*) indicate patients whose whole exomes were sequenced. (B) Patient’s details and hematological parameters. Hematological parameters of patients II.1 and II.2. MPV values are shown in the table as ‘Large’ because platelets with large volume are undetectable by the Sysmex analyzer. (C) The nonsense variant c.975T>G; p. Tyr325Ter results in the substitution of tyrosine residue at position 325 to a stop codon which is predicted to truncate the full length of 472 amino acids. (D) Mutation details (E) Western blot of protein from lysate from patient platelets showing truncation of the CD36 protein and GAPDH housekeeping loading control (F) Conservation of the tyrosine 325 residue across multiple species. The location of the tyrosine residue is shown by the highlighted green box. (G) Modelled structure of the CD36 ectodomain.^22^ The crystal structure shows the result of the CD36 nonsense variant on the structure of the WT CD36 protein (left panel) and mutant CD36 (right panel) as a result of the truncation. (H) Schematic of the CD36 protein structure. CD36 has two short cytoplasmic domains representing the C-terminal and N-terminals, two transmembrane domains and two large extracellular domains. The extracellular domain contains three disulfide bonds, binding sites of interaction with thrombospondin type I repeat (TSR), plasmodium falciparum, oxLDL, sites of acetylation (palmitoylation), phosphorylation, glycosylation, and the position of the nonsense variant found in patients II.1 and II.2.^57, 58^ (I) Protein expression of transfected CD36-wild type and CD36 mutants. C: pEF6 empty vector; WT: pEF6/CD36 wild type; Del: pEF6/CD36 deleted mutant; Sub: pEF6/CD36 substitution mutant. SDS-PAGE immunoblot expression analysis of samples probed with anti-CD36 and anti-GAPDH antibodies. Expected sizes of the samples are indicated on the right. (J) NFAT-luciferase activity measuring activation of CD36 after normalization of the stimulated and unstimulated conditions. Only WT CD36 shows luciferase activity over background. n=3 *M: male, F: female, WBC: white blood cell, RBC: red blood cell, Mono: monocyte, IPF: immature platelet fraction, MPV: mean platelet volume. WT: wildtype, Del: deletion, Sub: substitution*

**Table 1.** Patient details and hematologic parameters. Haematologic parameters of patients II.1 and II.2. M: male, F: female, WBC: white blood cell, RBC: red blood cell, Mono: monocyte, IPF: immature platelet fraction, MPV: mean platelet volume. MPV values were shown in the table as (Large) because platelets with large volume are undetectable by the Sysmex analyser. The Sysmex blood cell analyser showed the hematological parameters normal ranges which were taken from ^56^.

| **Patient** | **II.1** | **II.2** | **Normal range** |
| --- | --- | --- | --- |
| **Age** | 12-16 | 12-16 | — |
| **Sex** | M | F | — |
| **WBC** | 3.55 | 5.01 | M 3.91 – 10.90 x10<sup>9</sup>/L<br>F 4.49 – 12.68 |
| **RBC** | 4.73 | 3.87 | M 4.44 – 5.61 x10<sup>12</sup>/L<br>F 3.92 – 5.08 |
| **Monocytes** | 0.28 | 0.32 | M 0.29 – 0.95 x10<sup>9</sup>/L<br>F 0.25 – 0.84 |
| **Platelets** | 79 | 87 | M 166 – 308 x10<sup>9</sup>/L<br>F 173 - 390 |
| **IPF** | 24.4 | 36.8 | M 0.8 – 6.3 %<br>F 0.8 – 6.2 |
| **MPV** | Large | Large | M 9.3 – 12.1 fL<br>F 9.1 – 11.9 |

This variant has previously been reported at a relatively high frequency (0.08929) in the Afro-Caribbean population compared to the overall population (8.33e-3) and linked to thrombocytopenia^21^, however it has not been functionally characterized. The c.975T>G; p. Tyr325Ter variant was predicted to encode a truncated CD36 protein which lacked the carboxyl-terminal transmembrane domain and potential function of the CD36 protein. Fittingly, western blotting analysis using platelet lysates from the CD36-mutation positive patients revealed a truncated band (Fig. 6E). Of note, the tyrosine residue at position 325 is highly conserved across multiple species (Fig. 6F). We modelled the known crystal structure of the CD36 ectodomain based on Fu-Lien et al.^22^, showing the absence of a significant portion of the WT CD36 protein as a result of the CD36 nonsense variant, leading to the truncation and significant loss of domains crucial to the normal functioning of CD36 (Fig. 6G-H). Together, these modeling and experimental data support the conclusion that the c.975T>G; p. Tyr325Ter mutation leads to a truncation of the CD36 protein.

We further investigated the function of the CD36 mutant by generating constructs and cloning the WT CD36 cDNA into a pEF-BOS expression vector followed by site-directed mutagenesis to generate 2 mutant forms of CD36: (i) a deletion of the amino acids following the nonsense variant (Tyr325Ter) to the stop codon of the mature protein at amino acid residue 472 and (ii) a substitution of the point mutation only (c.975T>G, p.Tyr325Ter). Initially, the expression of the 3 constructs was measured and validated using western blotting, confirming the truncation effect of the 2 mutant CD36 constructs in both Jurkat T cells and HEK293 cells (Fig. 6I). Flow cytometry was then used to investigate the expression of CD36 mutant constructs on the cell surface. Of note, only WT CD36 and not the mutant constructs, was detected (Supplementary Fig. S5A), suggesting the other constructs were not trafficked to the cell surface. Next, we studied the signaling capacity of the CD36 constructs using a nuclear factor of activated T cells (NFAT)-luciferase reporter assay. The CD36 WT construct robustly activated the NFAT-luciferase, while the mutant constructs did not, confirming the absence of signaling capacity for the CD36 mutants (Fig. 6J). These data are consistent with the flow cytometry data and demonstrate that the mutant CD36 indeed does not reach the cell surface and does not signal, conferring a complete loss of function in the reported mutation. Together, our data reveal that in both humans and mice, loss of CD36 function resulted in reduced platelet counts, underscoring its importance in efficient platelet production.

## Discussion

Through lipidomics, we uncovered an enrichment in PUFAs, essential fatty acids that are primarily diet-derived, throughout MK differentiation and maturation and platelet production. Our data then revealed that fatty acid uptake, functionalization, and metabolism play differing but essential roles in MK differentiation and proplatelet production. Specifically, platelet production from MKs can be modified both in vitro and in vivo by varying the availability of exogenous PUFAs. We further demonstrated that CD36 is a key receptor responsible for the uptake of fatty acids in MK progenitors. We further report that a familial loss-of-function mutation in CD36 results in thrombocytopenia, suggesting that lipid uptake plays a critical role in platelet production.

Lipids are a vast class of biomolecules which fulfill three general functions: energy, membrane structure, and signaling.^4, 5^ To date, an impressive amount of experimental data has given insights into membrane biogenesis as well as homeostasis and lipid-protein interactions, which paves the way for targeted modification of membrane lipid compositions.^23^ In 1987, Dio et al. showed that adding SFAs (palmitic acid, etc.) to a murine fibroblast cell line results in a severe inhibition of cell growth.^24^ In contrast, increasing the PUFA concentration in liposomes results in a more flexible plasma membrane with a higher deformation rate in response to applied force.^25^ The idea of enhancing membrane flexibility by modulating PUFA content was further substantiated by Manni et al.^16^ Their study reveals that increasing the PUFA content in liposomes results in a higher tubulation rate. This process bears a striking resemblance to proplatelet production, which requires reorganization of the MK membrane system. Our data showed that increasing the dietary SFA:PUFA ratio resulted in a reduction in platelet generation both in vitro and in vivo. One way through which increased SFAs may be reducing proplatelet production is by creating a membrane that is too rigid to accommodate proplatelet extension. Conversely, replacing SFAs with PUFAs may ensure that the MK membrane is sufficiently flexible to allow for both DMS formation and proplatelet production. This is consistent with our lipidomic analysis revealing that MKs and platelets were increasingly enriched in PUFAs. These results highlight the potential to manipulate dietary fatty acid ratios and thereby modify MK phenotype and increase or decrease platelet production in vivo. Ultimately, these approaches may be able to fill a significant unmet clinical need in providing TPO-independent ways to modulate thrombopoiesis and mitigate abnormal platelet counts.

Changes in exogenous lipid availability and content are associated with a variety of diseases including obesity, which has become an alarming health problem worldwide.^26^ Previous studies have demonstrated that alterations in lipid metabolism and changes in plasma lipid profile are associated with the onset and progression of obesity-related complications.^27^ Often, however, obesity is associated with a pro-inflammatory and/or pro-thrombotic state that enhances the risk of developing cardiovascular diseases, where platelets play a well-established pathogenic role.^28–30^ In these settings, it is challenging to study the impact of obesity and an altered plasma lipid profile on MKs and platelets independent of comorbidities such as inflammation.^31, 32^ Recently, a new subset of obese individuals classified as ‘metabolically healthy obese’ (MHO) were found to be protected against worsening metabolic health.^33, 34^ Despite the debate about the use and clinical implications of MHO as a diagnosis, obesity without cardiometabolic abnormalities may provide a unique human model system to study mechanisms linking different diets and fat accumulation to obesity-related cardiometabolic complications.^33^ To date, no studies have reported how MHO may impact megakaryopoiesis or platelet production in humans. However, our in vitro data suggest that changes in plasma lipid content alone may lead to profound changes in MK maturation and platelet production, even in the absence of inflammation or other comorbidities associated with obesity. Notably, depending on the lipid profile, our data suggest that these changes may not always be pathogenic. Further, these data set the stage for future studies examining how changes in dietary lipids might also modify platelet function and reactivity.

CD36 is a multifunctional protein; one of its roles is to accelerate exogenous fatty acid uptake and incorporation into more complex lipids.^35, 36^ Here, we substantiated CD36 as a key receptor for the uptake of fatty acids in MKs and their progenitors and revealed that loss of CD36 led to thrombocytopenia in both mice and humans. These findings align with recent studies that demonstrate fatty acid uptake transfer between adipocytes and MKs is dependent on CD36.^17^ Notably, evidence suggests that there is little expression of CD36 on HSCs, but expression increases dramatically in MKs^37^, providing a possible mechanism for how they rapidly accumulate fatty acids over maturation via a cell intrinsic manner.

There are also important limitations to this work that should be noted. First, we only identified and characterized two patients with the c.975T>G, p.Tyr325Ter CD36 mutation. However, our data are consistent with previous studies that also linked this and similar mutations to thrombocytopenia.^21^ In addition, the fact that *Cd36^−/−^* mice and humans with mutated or absent CD36 have both MKs and platelets, albeit at reduced levels, suggests that other receptors, such as the free fatty acid receptors (FFARs) are also involved in fatty acid uptake.^38^ Specifically, FFAR_2_ may be a plausible candidate as it is expressed in MKs and their precursor cells.^39^

In summary, our data provide unique insights into the functional role of lipids, diet, and lipid metabolism during MK differentiation, maturation, and platelet production. We have further validated an essential receptor, CD36, as a key mechanism for lipid uptake and identified a CD36 mutation that is a genetic determinant of thrombocytopenia and pathological bleeding. In the future, we aim to identify additional MK-specific lipid signatures to determine novel ways to increase or decrease megakaryopoiesis and platelet production. This could be as simple as altering fatty acid ratios in the diet or involve therapeutics that target key fatty acid biosynthesis enzymes or the uptake receptor CD36. Ultimately these approaches may be able to meet a significant unmet clinical need in the modulation of thrombopoiesis to mitigate abnormal platelet counts.

## Methods

### Animal Models

CD-1 and C57BL/6J mice were acquired from Charles River Laboratories (Worcester, MA) or The Jackson Laboratory (Bar Harbor, ME). Mice were housed in the animal facilities at Boston Children’s Hospital, Boston, MA or Michigan State University, East Lansing, MI. All animal work was approved by the international animal care and use committee at Boston Children’s Hospital, Boston, MA (00001248, and 00001423) and Michigan State University (PROTO201800186).

For experiments with high fat diets, adult C57BL/6J and *Cd36^−/−^* mice were placed on the following diets: 1) Research diets Inc. D12492, Rodent Diet With 60 kcal% from Fat and 2) Research diets Inc. D22050406i, Rodent Diet with 60 Kcal% Fat enriched in polyunsaturated fatty acids. As a control, mice were fed a chow diet (D12450B, Research diets Inc.) Mice received water and food ad libitum. Mice were sacrificed at the indicated endpoints using methods consistent with IACUC protocol guidelines. CD36 knockout mice (*Cd36^−/−^*) were obtained from The Jackson Laboratory (Strain #019006). Homozygous mice were bred to maintain a colony of knockout mice. Ear clipping specimens were sent to Transnetyx, Inc for genotyping to ensure integrity of the strain and preservation of the *Cd36* gene knockout.

### Blood parameters

For non-terminal blood collection, mice were anesthetized with isoflurane and of 70 µL of blood collected from the retro-orbital capillary bed using heparinized capillaries, and transferred into EDTA-coated tubes (BD Microtainer™ Capillary Blood Collector). Platelet count, size, and basic blood parameters such as white and red blood cell counts were obtained using an automated cell counter (XN-1000™ Sysmex).

### Thiazole Orange analysis

Whole blood (5 µL) was diluted in 500 µL Tyrode’s buffer. 50 µL of diluted blood was incubated with 200 ng mL^-1^ of thiazole orange (TO) and an antibody against CD41/61 (Emfret Analytics) was used to identify platelets. TO-positive platelets were identified by flow cytometry (Accuri C6 plus, BD Biosciences).

### Platelet Isolation

For terminal bleeds, mice were anaesthetized using 2% isoflurane and blood was collected from the inferior vena cava into EDTA containing tubes. Platelet-rich plasma (PRP) was obtained by centrifugation at 200*g* for 10 minutes (acceleration 9, break 6). After addition of PGI1 (final concentration 560 nM, P5515 Sigma) and Apyrase (final concentration 0.02 U/mL, A6123-500U Sigma), PRP was spun down at 800*g* for 10 minutes (acceleration 9, brake 6). Platelets were washed twice in modified Tyrode’s buffer (134 mM NaCl, 2.9 mM KCl, 20 mM HEPES, 1 mM MgCl_2_, 5 mM Glucose, pH 6.5) containing PGI1 and Apyrase (centrifugation for 10 min at 800*g* with acceleration 9 and break 6). After the last wash, the platelet pellet was resuspended in modified Tyrode’s buffer. Platelets were counted on the ProCyte Dx and diluted in PBS to a final concentration of 10,000 platelets/µL. Samples were stored at −80°C prior to lipidomics studies.

### Isolation of murine fetal liver MKs

Fetal liver MKs were derived from fetal livers that were cultured as previously described.^40^ Briefly, CD-1 pregnant mice at day 13.5 of gestation mice were sacrificed by CO_2_ asphyxiation followed by cervical dislocation and fetal livers were extracted. Homogenized fetal liver cells were then cultured in complete media (Dulbecco’s Modified Eagle Medium (Sigma), 10% Fetal Bovine Serum, (Sigma) and 1% Penicillin Streptomycin (Gibco) in the presence of recombinant murine thrombopoietin (TPO; 50 ng/mL) for 4 days. On day 4, mature MKs were enriched using a bovine serum albumin (BSA) density gradient.

### Isolation of murine bone marrow MKs

Mice were sacrificed using methods consistent with IACUC protocol guidelines. Long bones and iliac crests were isolated and bone marrow was obtained by centrifugation at 2500*g* for 40s as previously described.^41^ Hematopoietic stem and progenitor cells (HSPCs) were separated by lineage depletion using an antibody mixture (Lineage depletion panel, 133307, Biolegend) and magnetic beads (CD4 untouched, 11415D, Invitrogen). For MK maturation, HSPCs were incubated in complete medium containing TPO (50 ng/mL) and recombinant hirudin (100 U/mL, Aniara Diagnostics, RE120A)) for 4 days.^42^ To assess proplatelet production, differentiated MKs were enriched by a BSA density gradient and used either directly or further incubated in the presence hirudin for 24h. Cells were imaged on a microscope (Nikon Eclipse Ts2) and mean cell size was measured using ImageJ Software (NIH).

### HSPC panel

Mice were sacrificed using methods consistent with IACUC protocol guidelines. Long bones and iliac crests were isolated, and bone marrow was obtained by centrifugation as described above. Bone marrow cells were filtered using a 70 μm PluriStrainer filter (Pluriselect). Red blood cells lysis was performed according to the manufacturer’s instructions (Lysis Buffer, 555899, BD Biosciences). Cells were stained as follows:

| Antibody | Fluorophore | Manufacturer | Catalog |
| --- | --- | --- | --- |
| Lin cocktail | PB | Biolegend | 133305 |
| cKit | APC | eBioscience | 17-1171-82 |
| SCA | PERCPCy5.5 | Biolegend | 108123 |
| CD41 | BV605 | Biolegend | 133921 |
| CD48 | APCy7 | Biolegend | 103432 |
| CD34 | BV786 | BD | 742971 |
| CD150 | PECy5 | Biolegend | 115912 |
| CD13 | PE | BD | 558745 |
| CD105 | PECy7 | Biolegend | 120409 |

For each sample, 1 μL of each antibody (1:100) and 50 μL of BV staining buffer was added. Unstained samples from both groups, single stained samples, single color beads (1 μL of antibody and 1 drop of UltraComp eBeads Plus Compensation Beads, Invitrogen), and FMO controls were were used. Cells were stained for 30 min, followed by 2 consecutive washes with MACS running buffer (Miltenyi). Prior to measuring the sample, DAPI (150 nM) was added, and cells were filtered using a 70 μm PluriStrainer filter. Samples were analyzed using a Spectral Flow Cytometry, Cytek Aurora (Cytekbio).

### Cell sorting for lipidomics

Mice were sacrificed using methods consistent with IACUC protocol guidelines. Long bones and iliac crests were isolated, and bone marrow was obtained by centrifugation as described above. Bone marrow cells were filtered using a 100 μm filter (Pluriselect). Red blood cell lysis was performed according to the manufacturers’ instructions (Lysis Buffer, 555899, BD Biosciences). MEP (Lin-, Sca-1, c-Kit+, CD34-, FcgR-, 10,000 cells) and MK (immature: CD41+, and mature: CD41+, CD42d+ double positive, 10,000 cells) populations were isolated using fluorescence activated cell sorting according to published studies.^43^ Samples were sorted using a BD FACS Aria IIu (BD bioscience).

### Lipid extraction

Cell samples were lyophilized using either a Savant SpeedVac (Thermo Scientific) or a CoolSafe freeze dryer (ScanVac) prior to extraction and resuspended in 10 µL MilliQ H_2_O. Lipids were extracted using a modified single-phase chloroform/methanol extraction method^44^. Briefly, 200 µL chloroform-methanol (2:1) was added to each sample along with an internal standard (ITSD) mixture containing stable-isotope labelled or non-physiological lipids. In tandem, blank control samples and plasma QCs were extracted and dispersed evenly throughout the extraction order to ensure optimal assay performance and to monitor variation that may arise from the extraction. Samples were subsequently mixed with a rotary mixer for 10 minutes at 90 rpm, sonicated for 30 mins at room temperature and centrifuged at 13,000 rpm for 10 mins to precipitate proteins from the lipid extracts. Supernatant containing the extracted lipids were transferred to a 96 well plate and evaporated using a Savant SpeedVac. Once dried, extracts were reconstituted in H_2_O-saturated butanol and methanol with 10 mM ammonium formate and moved to glass vials and stored until mass spectrometry analysis.

### Liquid chromatography tandem mass spectrometry (LC-MS/MS)

Lipid extracts were analyzed using an Agilent 6490 triple quadrupole (QqQ) mass spectrometer coupled to an Agilent 1290 high performance liquid chromatography (HPLC) system as previously published^45^. Briefly, we used a ZORBAX eclipse plus C18 column (2.1×100mm 1.8µm, Agilent) with thermostat set to 45°C. The final mass spectrometry analysis on each cell population was performed in positive mode with dynamic scheduled MRM. Solvents consisted of solvent A (50% H2O, 30% acetonitrile, 20% isopropanol with 10 mM ammonium formate) and solvent B (1% H2O, 9% acetonitrile, 90% isopropanol with 10 mM ammonium formate) and followed a modified 20-minute gradient as follows:

A wash vial comprising of 1:1 butanol:methanol was used after each sample injection. To further improve chromatic peak shape for many anionic/acidic lipid species (notably PS, PA, PIP, S1P), an additional pre-run passivation step was done with phosphoric acid to minimize interaction between the HPLC unit and these lipids.

| Time (min) | Solvent A (%) | Solvent B (%) | Flow (mL/min) |
| --- | --- | --- | --- |
| 0.00 | 85.00 | 15.00 | 0.4 |
| 2.50 | 48.00 | 52.00 | 0.4 |
| 10.00 | 40.00 | 60.00 | 0.4 |
| 13.00 | 15.00 | 85.00 | 0.4 |
| 17.00 | 7.00 | 93.00 | 0.4 |
| 17.10 | 0.00 | 100.00 | 0.4 |
| 17.90 | 0.00 | 100.00 | 0.4 |
| 18.00 | 85.00 | 15.00 | 0.4 |
| 20.00 | 85.00 | 15.00 | 0.4 |

### Lipid nomenclature

The lipid names used follow guidelines set by the LIPIDMAPS consortium.^46^ Phospholipids with detailed characteristics i.e., acyl chain composition are annotated as [PC(16:0_20:4)] with PC being the lipid class and (16:0_20:4) representing the acyl chains found on the glycerol backbone, irrespective of sn1 or sn2 position. Lipids without specific structural annotations are named based on their sum acyl chain length and degrees of saturation e.g. PC (36:4). Isomeric lipid species separated chromatographically but incompletely annotated were designated (a), (b) etc., with (a) and (b) representing elution order. Owing to technical limitations, we were unable to assign acyl chains to a specific sn1 or sn2 position for the majority of PL species, only that the acyl chain was present at either the sn1 or sn2 position. In the case of ether-PC and PE, the alkyl or alkenyl chains were always located at the sn1 position and the acyl chain always at the sn2 position. In addition, we were unable to determine acyl chain composition for all PLs and the data shown therefore represents those PLs for which we were able to obtain information on their structural composition. Notably, this represents >90% of total PLs for all cell types.

### Analysis of Lipids

Individual analyte areas were divided by the area of the corresponding internal standards (ITSDs), and the median of ITSD containing blank samples was subtracted from each analyte (background subtraction). Background subtracted values were multiplied by the concentration of the ITSD and the analyte’s respective response factor (Rf). Zeroed background subtraction values (i.e., sample values which were lower than the median blank + ITSD samples) were replaced with 1/10th of the minimum value for the corresponding analyte. Data was ultimately normalized to pmol/µmol total lipidome where background subtracted data for an individual lipid was divided by the sum of the total lipidome of the sample and multiplied by a factor of 10^6^. Normalized raw data can be found in supplemental data.

### Lipid Enrichment Analysis

Lipid ontology analysis was performed using Lipid Ontology (LION). Analysis was conducted in ranking mode with lipidomic data normalized to mol%. LION-term enrichment was considered significant when FDR q value < 0.05.

### RNAseq dataset and Enrichment Analysis

The mRNA seq dataset was previously published by our group.^19^ To corroborate if lipid-related genes were differently regulated in non-proplatelet-forming MKs vs proplatelet-forming MKs, the dataset was examined for keywords (e.g lipid, phospholipid, coA, fatty acid, CD36). Selected genes with p value and fold change were plotted using Prisma. Enrichment analysis was done using Reactome.^47^

### Flow cytometry on mature MKs

At day 4, mature MKs (bone marrow or fetal liver) were centrifuged and washed with autoMACS® Running Buffer (130-091-221, Miltenyi). Cells were stained for 30 min using CD41-FITC (133904, BioLegend) and CD42d-APC (148506, BioLegend) antibodies. Unstained and single cell controls were included. After the incubation, cells were washed twice and analyzed on a FACScalibur (BD Biosciences). The percentage of CD41- and CD41/CD42d-positive cells was analyzed using FlowJo and percentage were normalized to the vehicle.

### Cytotoxicity assay

The lactate dehydrogenase (LDH) cytotoxicity assay was performed according to the manufacturers’ instructions (C20300, ThermoFisher Scientific). Mature FLMKs were treated with indicated dosages of Triacsin C (T4540, Sigma), PF-05175175 (PZ0299, Sigma), and Cerulenin (C2389, Sigma) and incubated overnight in a 96-well-plate and LDH activity in the supernatant was measured. As a positive control, cells were lysed with TritonX-100. MKs treated with vehicle were treated as a negative control (baseline).

### Proplatelet formation using the Incucyte automated microscope

Fetal liver- or bone marrow-derived MKs were isolated as described above. Immediately following density gradient enrichment, MKs were either untreated, supplemented with different fatty acids, or treated with inhibitors (Triacsin C, T4540, Sigma), PF-05175175 (PZ0299, Sigma), Cerulenin (C2389, Sigma), as indicated. For the supplementation, MK were treated with palmitic acid (P0500, Sigma) or arachidonic acid (A-122, Sigma) in serum-free media. Proplatelet formation was visualized on an Incucyte imaging system and quantified using a custom image analysis pipeline or manually by counting the percentage of MKs making proplatelets, using Image J.^48^

### Immunostaining of proplatelet-forming MKs

μ-side 8-well Ibidi chambers were coated with an anti-CD31 antibody (102502, BioLegend) for 30 min, followed by blocking with 3% BSA for 30 min. Bone marrow derived MKs were isolated as described above. Immediately following density gradient enrichment, MKs were treated with the inhibitors or supplemented with the different fatty acids (see below). After treatment, cells were pipetted gently onto the bottom of the chamber and incubated overnight. The following day, cells were fixed using 4% PFA containing 0.1% Tween20 for 30 min, followed by 3% BSA, and stained for α-tubulin (A488, 322588, ThermoFisher), F-actin (Phalloidin-Atto647N, A22287), and DAPI (Sigma) overnight. Proplatelet formation was visualized using a Zeiss LSM880 confocal microscope (40x objective).

### Oxygen Consumption Measurement

MKs were suspended in XF base medium DMEM (Agilent Bioscience) supplemented with 1 mM sodium pyruvate (Agilent Bioscience), 2 mM glutamine (Wisent), and 10 mM glucose (Agilent Bioscience), pH 7.4. A total of 10,000 MKs per well was seeded on XF-96 plates (Agilent/Seahorse Bioscience). Cells were treated with Triacsin C (T4540, Sigma), PF-05175175 (PZ0299, Sigma), Cerulenin (C2389, Sigma) at the indicated concentrations in complete media for 90 min prior the measurement. Plates were then centrifugated at 300*g* for 2 min at room temperature. We ensured cell homogeneous repartition under a microscope and plates were maintained at 37°C without CO_2_ for approximately 60 min prior to loading. Oxygen consumption rates were measured in accordance with manufacturer instructions (Agilent/Seahorse Bioscience). Experiments were replicated in three to five wells and averaged for each experimental condition. A total of 3 measurements of oxygen consumption for each condition were made approximately every 10 min (mix for 3 min, wait for 4 min and measure for 3 min) under basal conditions and after sequential injection of oligomycin (4 μM), FCCP (carbonyl cyanide 4-(trifluoromethoxy) phenylhydrazone, (1 μM) and rotenone/antimycin A (1 μM each). Oligomycin is used as an ATP synthase inhibitor, FCCP as an uncoupling agent of oxidative phosphorylation, rotenone as a complex I inhibitor and antimycin A as a complex III inhibitor. This allowed us to estimate the contribution of individual parameters for basal respiration, proton leak, maximal respiration, spare respiratory capacity, non-mitochondrial respiration, and ATP production.

### Click-chemistry

HSPCs were isolated as described above. After lineage depletion, cells were supplemented with fatty acids modified with a ω-terminal alkyne group (Palmitic acid, 13266; Arachidonic Acid, 10538; Cayman Chemical). After 4 days of maturation, click-chemistry was performed using green-fluorescent Alexa Fluor® 488 azide (C10641, ThermoFisher Scientific) according to the manufacturers’ instructions. Azides are specifically reactive with terminal alkynes via a copper-catalyzed click reaction.^49, 50^ Cells were image using Zeiss LSM880 confocal microscope (20x and 63x objectives). For the analysis, Alexa Fluor® 488 MFI was measured using Image J and normalized to the vehicle.

### MK ploidy

Bone marrow was isolated from 1 femur by centrifugation at 2500*g* for 40 seconds. Cells were filtered through a 100 µm cell strainer and red blood cells were lysed using ACK buffer (A1049201, Gibco). Cells were washed in PBS, fixed, and permeabilized in 100% ethanol for 30 min on ice. Cells were treated with RNAse A (EN0531, ThermoFisher Scientific) and stained with an an ti-CD41-FITC antibody (133904, BioLegend) and propidium iodide (P1304-MP, Sigma Aldrich) for 30 min on ice. Ploidy distribution and percentage of CD41-positive cells were quantified by flow cytometry (BD AccuriC6 Plus).

### Cryosectioning and immunofluorescence staining

Mice were sacrificed and femurs were isolated and fixed in 4% PFA overnight. Femurs were transferred into 10% sucrose in PBS and a sucrose gradient was performed over 3 days. Femurs were sectioned at 10 μm using the Cryostat CM3050 S (Leica Biosystem), transferred onto slides using a tape-transfer system^51^, and rehydrated in PBS for 15 min. Sections were blocked using 5% goat serum and stained using antibodies against CD41 (133902, Biolegend) and laminin (L9393, Sigma) overnight. The following day, sections were incubated with secondary antibodies (goat anti-rat A488 and goat anti-rabbit A647, respectively). After washing in PBS containing 0.1% TritonX 100, DAPI was added for 5 min and the slides were mounted using Fluoroshield (Sigma Aldrich). Image acquisition was performed at a confocal microscope using a 20x objective (Zeiss LSM880,). MK numbers and area were quantified manually using ImageJ Software (NIH).

### Patient recruitment and testing

Patients were consented and recruited to the GAPP study from multiple collaborating Hemophilia Centers across the UK and Ireland as previously described^52^ and approved by the UK National Research Ethics Service by the Research Ethics Committee of West Midlands (06/MRE07/36). The study cohort currently consists of >1000 patients with a history of bleeding and suspected of having a platelet disorder of unknown cause. Platelet counts, mean platelet volume (MPV) and other hematological parameters were measured on the Sysmex Whole Blood Analyzer. Whole exome sequencing (WES) was performed in patient genomic DNA as previously reported.^53^ For WES filtering of candidate genetic variants was performed to identify rare variants and final sequence variants were confirmed in patients using Sanger sequencing.

### Functional analysis of CD36 mutants and expression

The Q5 Site-Directed Mutagenesis (SDM) Kit (NEB®, USA, #E0554S) was used to introduce the *CD36* heterozygous stop gain variant (c.975T>G; p. Tyr325Ter) into the human *CD36* cDNA wild-type cloned into the mammalian expression pEF-BOS^54^ using the following primers:

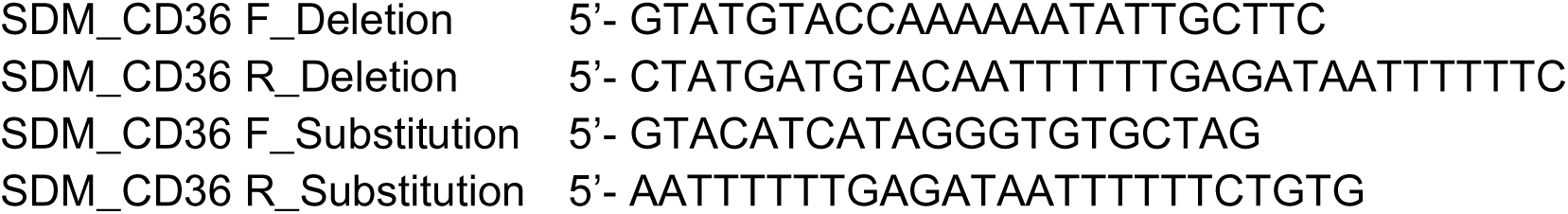

The NFAT-luciferase reporter assay was used as previously described.^55^ Briefly, Jurkat T cells were electroporated with 20 µg NFAT-luciferase construct and 12.5 µg of either WT CD36 or mutant CD36 constructs. Twenty-four hours post transfection, cells were harvested and purified, and flow cytometry and western blot analyses were used to assess expression levels.

### Data analyses

The presented results are mean ± standard deviation (SD). Data distribution was analyzed using the Shapiro-Wilk-test and differences between control and mutant mice were statistically analyzed using unpaired, two-tailed Student’s t-test, and one- or two-way ANOVA. Tukey or Sidak’s post-hoc tests were used for multiple comparisons, as indicated. P-values < 0.05 were considered statistically significant.

## Acknowledgements

KRM is supported by grants from the National Institutes of Health, National Institute of Diabetes and Digestive and Kidney Diseases (R03DK124746) and National Heart, Lung, and Blood Institute (R01HL151494). MNB is supported by fellowships from the American Society of Hematology and the American Heart Foundation. ICB is supported by a Walter Benjamin Fellowship of the German Research Foundation (DFG; BE 7766/2-1). TH is supported by NIH grant 5T32HL007734 and The Boston Children’s Hospital Surgical Foundation. MP is supported by The Boston Children’s Hospital Surgical Foundation. AOK is a Sir Henry Wellcome Fellow supported by the Wellcome Trust (218649/Z/19/Z). IA received support from the Saudi Arabia Cultural Bureau in London. AJM is supported by an NHMRC investigator grant (APP1194329) and a CSL Centenary Award. JEI is supported by the National Institute of Health National Heart, Lung and Blood Institute (R35HL161175). EB is recipient of a senior award from the Fonds de Recherche en Santé du Québec. The work was performed in part through the support of a Project grant from the Canadian Institutes of Health Research. JPL is supported by grants from the National Institutes of Health, National Institute of Diabetes and Digestive and Kidney Diseases (R01DK112778) and support from the US Department of Agriculture (USDA) National Institute of Food and Agriculture (to JPL).

We would like to thank Drs. Connie Koo and Mike Tomlinson for assistance with *in vitro* characterization of the CD36 mutation and Dr Beatrice Nolan for referring the patients to the GAPP study. We would like to thank Dr. Mathieu Laplante and his team for their help with in vivo experiments and for being an excellent collaborator. Thank you to Dr. Jessica Cardenas for her critical review of the manuscript and excellent edits and to D/Z Rotfus and JJ Freire for their unwavering patience and support.

## Author Contributions

MNB conceived the study, performed experiments, collected, and analyzed data, and wrote the manuscript. GP, ICB and IA performed experiments, collected, and analyzed data. AOK provided intellectual input. E.C designed and developed image analysis methodologies. DJG, DF, KG, ZW and IA performed experiments and analyzed data.TH and MP maintained the CD36 knockout mouse colony. JPL performed experiments and analyzed data. NVM recruited and governed the patient’s ethics, performed experiments, analyzed data, and helped to draft the manuscript.. PJM and TJC analyzed data and prepared figures. NAM performed experiments and analyzed data. PJM and JEI provided intellectual input and key reagents. EB provided intellectual input and interpreted data. AJM conceived the study, interpreted the data and helped write the manuscript. KRM conceived and directed the study, analyzed data, and wrote the manuscript.

All authors provided input on and reviewed the manuscript.

## Ethics Declarations

### Competing Interests

All authors have no conflicts of interest to declare that are relevant to the content of this article.

## Supplementary Figure Legends

**Supplementary Figure 1. Megakaryocytes and platelets have unique lipidomic profiles.** Murine bone marrow cell populations were isolated by fluorescence-activated cell sorting and platelets by sequential centrifugation. Lipids were extracted and analyzed using 20-min gradient HPLC and mass spectrometry (see methods for details). (A) Percentage of different lipid classes of indicated murine bone marrow cell populations and autologous platelets in lipidomic analyses. Volcano plots from the different lipid classes between (B) bone marrow extracellular fluid (BMEF) and MEPs. n=4 and 8, respectively, (C) BMEF and immature (CD41+) MKs, n=4 (D) BMEF and mature (CD41/42+) MKs, n=4, and (E) platelets and plasma n=4. Total percentage of all acyl/alkyl composition of (F) PC (G) PI, (H) PE, (I) PS. n=8 for MEP and 4 for all other cell types.

*MK: megakaryocyte; MEP: MK-erythroid progenitor; PA: phosphatidic acid; PC: phosphatidylcholine; PE: phosphatidylethanolamine; PI: phosphatidylinositol; PS: phosphatidylserine; PG: phosphatidylglycerol*

**Supplementary Figure 2. Fatty acid uptake and synthesis affected MK differentiation but not mitochondria metabolism.** (A) Fetal liver-derived HSPCs were cultured with TPO and treated with the ACSL inhibitor Triacsin C at indicated doses. CD41+ and CD41/CD42d+ cells were quantified using flow cytometry, n=5, one-way ANOVA – Dunnett’s test. (B) Cytotoxicity assay was performed on day 4 after fetal liver MKs were treated with Triacsin C. Control=cells treated with the vehicle; positive control=cells lysed with TritonX-100. n=4. (C) Fetal liver derived HSPCs were cultured with TPO and treated with the ACC inhibitor PF-05175175 at indicated doses. CD41+ and CD41/CD42d+ cells were quantified using flow cytometry, n=5, one-way ANOVA – Dunnett’s test. (D) Cytotoxicity assay was performed day 4 after fetal liver MKs treated with PF-05175175. Control=cells treated with the vehicle; positive control=cells lysed with TritonX-100. n=4 (E) Fetal liver derived HSPCs were cultured with TPO and treated with the FASN inhibitor Cerulenin at indicated doses CD41+ and CD41/CD42d+ cells were quantified using flow cytometry, n=5, one-way ANOVA – Dunnett’s test. (F) Cytotoxicity assay was performed day 4 after fetal liver MKs were treated with Cerulenin. Control=cells treated with the vehicle; positive control=cells lysed with TritonX-100. n=4 (G) Representative graph of mitostress assay. MKs were treated with Triacsin C (T4540, Sigma), PF-05175175 (PZ0299, Sigma), Cerulenin (C2389, Sigma) at the indicated concentrations in complete media for 90 min prior the measurement. Oxygen consumption rates were measured in accordance with manufacturer instructions (Agilent/Seahorse Bioscience) (H) Individual parameters for basal respiration, maximal respiration, and spare respiratory capacity were measured and analyzed. n=5, one-way ANOVA.

**Supplementary Figure 3. Blood cell counts and characteristics in SFA-enriched high fat diet.** Blood parameters were measured using a Sysmex hematology analyzer, (A) Platelet Distribution Width (PDW), (B) Mean Platelet Volume (MPV), (C) red blood cells, (D) white blood cells, (E) neutrophils; n=12-16, unpaired t-test.

**Supplementary Figure 4. Blood cell counts and characteristics in PUFA-enriched high fat diet.** Blood parameters were measured using a Sysmex hematology analyzer, (A) Platelet Distribution Width (PDW), (B) Mean Platelet Volume (MPV), (C) red blood cells; n=10, unpaired t-test.

**Supplementary Figure 5. Mutant CD36 constructs are not trafficked to the cell surface.** Wildtype (WT) and mutant CD36 constructs were transfected into both A) Jurkat T cells (n=3) and B) HEK293 cells and surface expression was assessed. Flow cytometry scatter plots indicate that only wildtype CD36 is detected on the cell surface for both cell types.

## Notes

### Competing Interest Statement

The authors have declared no competing interest.

