## Supplemental Figures for "Efficient megakaryopoiesis and platelet production require phospholipid remodeling and PUFA uptake through CD36"

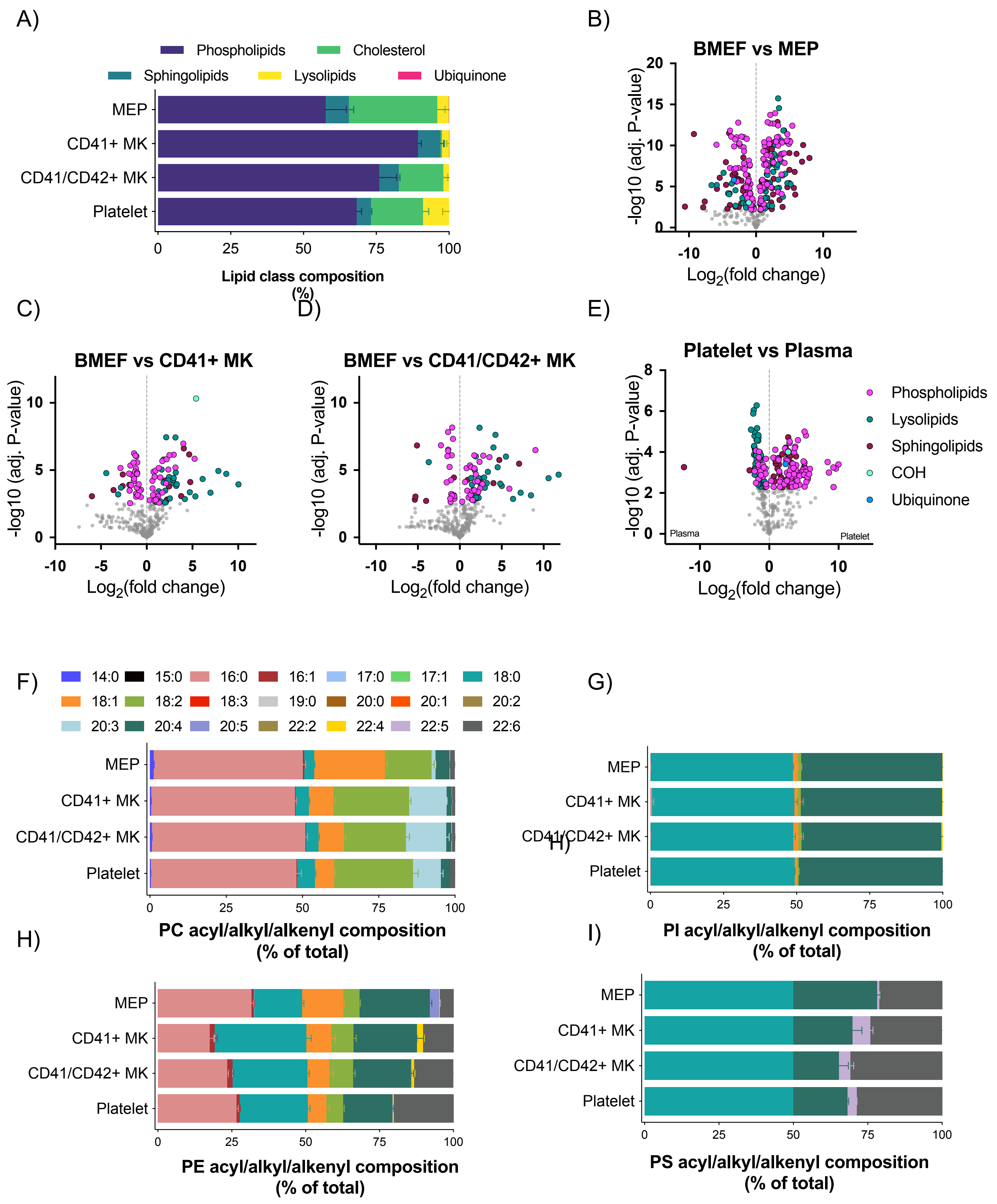


Supplementary Figure 1

**Supplementary Figure 1. Megakaryocytes and platelets have unique lipidomic profiles**. Murine bone marrow cell populations were isolated by fluorescence-activated cell sorting and platelets by sequential centrifugation. Lipids were extracted and analyzed using 20-min gradient HPLC and mass spectrometry (see methods for details). (A) Percentage of different lipid classes of indicated murine bone marrow cell populations and autologous platelets in lipidomic analyses. Volcano plots from the different lipid classes between (B) bone marrow extracellular fluid (BMEF) and MEPs. n=4 and 8, respectively, (C) BMEF and immature (CD41+) MKs, n=4 (D) BMEF and mature (CD41/42+) MKs, n=4, and (E) platelets and plasma n=4. Total percentage of all acyl/alkyl composition of (F) PC (G) PI, (H) PE, (I) PS. n=8 for MEP and 4 for all other cell types.

*MK: megakaryocyte; MEP: MK-erythroid progenitor; PA: phosphatidic acid; PC: phosphatidylcholine; PE: phosphatidylethanolamine; PI: phosphatidylinositol; PS: phosphatidylserine; PG: phosphatidylglycerol*

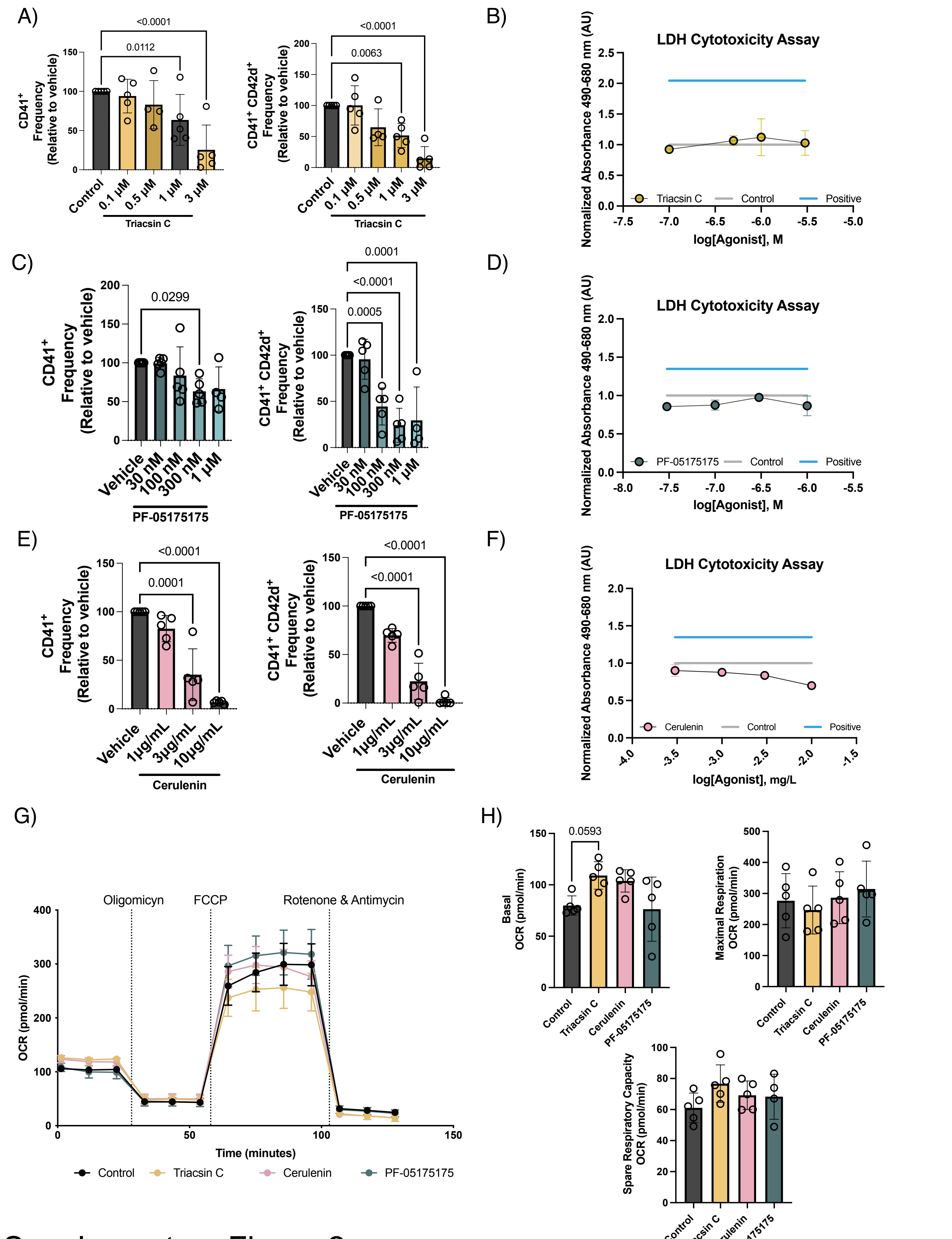


### Supplementary Figure 2

**Supplementary Figure 2. Fatty acid uptake and synthesis affected MK differentiation but not mitochondria metabolism**. (A) Fetal liver-derived HSPCs were cultured with TPO and treated with the ACSL inhibitor Triacsin C at indicated doses. CD41+ and CD41/CD42d+ cells were quantified using flow cytometry, n=5, one-way ANOVA – Dunnett's test. (B) Cytotoxicity assay was performed on day 4 after fetal liver MKs were treated with Triacsin C. Control=cells treated with the vehicle; positive control=cells lysed with TritonX-100. n=4. (C) Fetal liver derived HSPCs were cultured with TPO and treated with the ACC inhibitor PF-05175175 at indicated doses. CD41+ and CD41/CD42d+ cells were quantified using flow cytometry, n=5, one-way ANOVA – Dunnett's test. (D) Cytotoxicity assay was performed day 4 after fetal liver MKs treated with PF-05175175. Control=cells treated with the vehicle; positive control=cells lysed with TritonX-100. n=4 (E) Fetal liver derived HSPCs were cultured with TPO and treated with the FASN inhibitor Cerulenin at indicated doses CD41+ and CD41/CD42d+ cells were quantified using flow cytometry, n=5, one-way ANOVA – Dunnett's test. (F) Cytotoxicity assay was performed day 4 after fetal liver MKs were treated with Cerulenin. Control=cells treated with the vehicle; positive control=cells lysed with TritonX-100. n=4 (G) Representative graph of mitostress assay. MKs were treated with Triacsin C (T4540, Sigma), PF-05175175 (PZ0299, Sigma), Cerulenin (C2389, Sigma) at the indicated concentrations in complete media for 90 min prior the measurement. Oxygen consumption rates were measured in accordance with manufacturer instructions (Agilent/Seahorse Bioscience) (H) Individual parameters for basal respiration, maximal respiration, and spare respiratory capacity were measured and analyzed. n=5, one-way ANOVA.

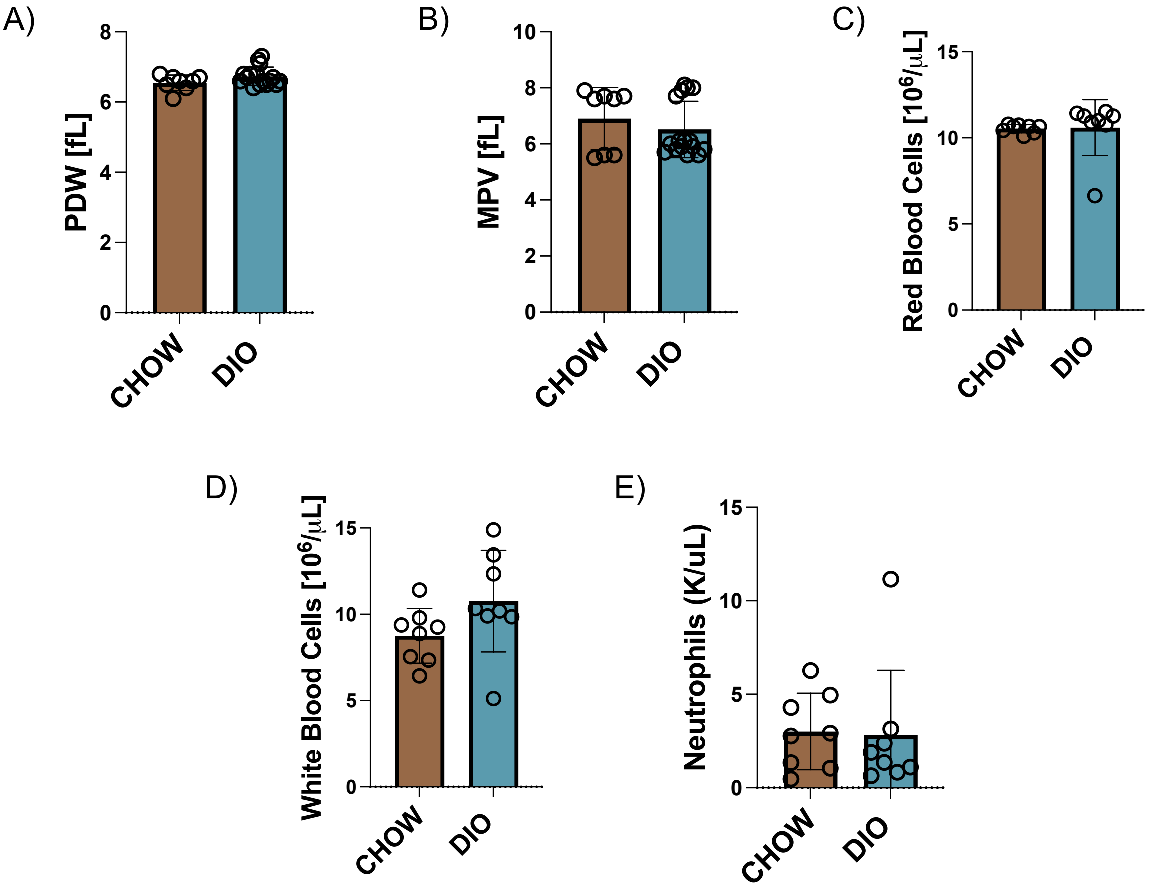


Supplementary Figure 3

**Supplementary Figure 3. Blood cell counts and characteristics in SFA-enriched high fat diet.** Blood parameters were measured using a Sysmex hematology analyzer, (A) Platelet Distribution Width (PDW), (B) Mean Platelet Volume (MPV), (C) red blood cells, (D) white blood cells, (E) neutrophils; n=12-16, unpaired t-test.

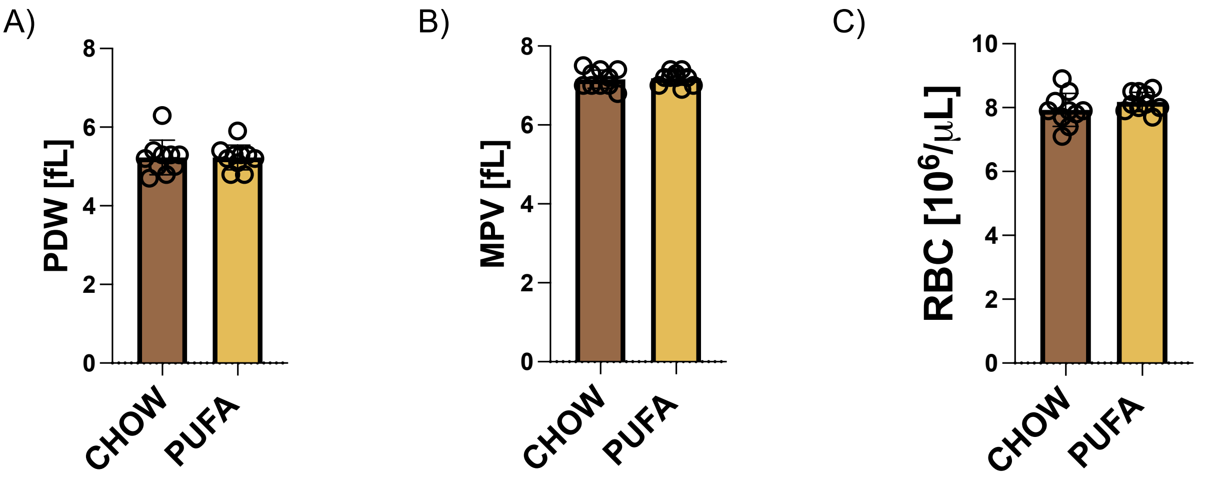


Supplementary Figure 4

**Supplementary Figure 4. Blood cell counts and characteristics in PUFA-enriched high fat diet.** Blood parameters were measured using a Sysmex hematology analyzer, (A) Platelet Distribution Width (PDW), (B) Mean Platelet Volume (MPV), (C) red blood cells; n=10, unpaired t-test.

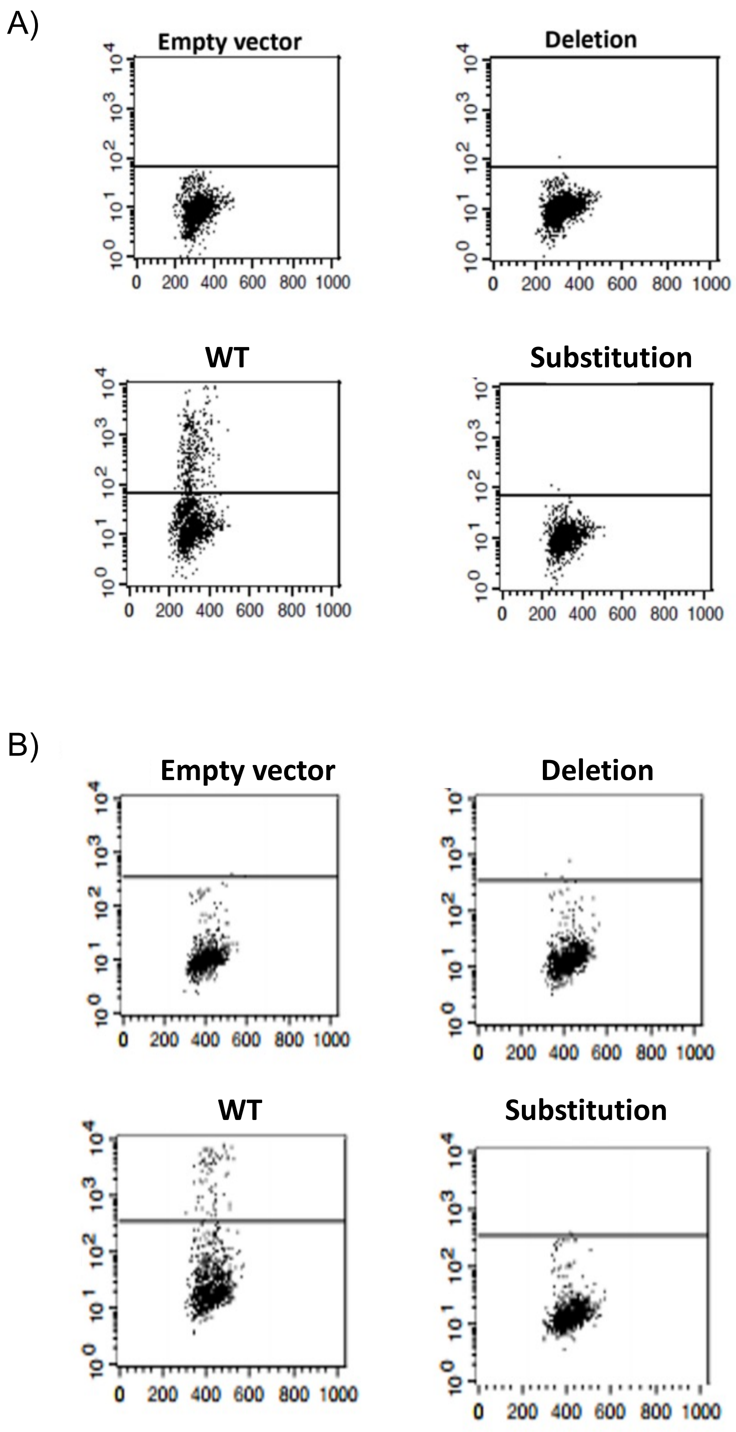


Supplementary Figure 5

**Supplementary Figure 5. Mutant CD36 constructs are not trafficked to the cell surface.** Wildtype (WT) and mutant CD36 constructs were transfected into both A) Jurkat T cells (n=3) and B) HEK293 cells and surface expression was assessed. Flow cytometry scatter plots indicate that only wildtype CD36 is detected on the cell surface for both cell types.
